# The sequestration of miR-642a-3p by a complex formed by HIV-1 Gag and human Dicer increases AFF4 expression and viral production

**DOI:** 10.1101/2023.05.24.542197

**Authors:** Sergio P. Alpuche-Lazcano, Owen R. S. Dunkley, Robert J. Scarborough, Sylvanne M. Daniels, Aïcha Daher, Marin Truchi, Mario C. Estable, Bernard Mari, Andrew J. Mouland, Anne Gatignol

## Abstract

Micro (mi)RNAs are critical regulators of gene expression in human cells, the functions of which can be affected during viral replication. Here, we show that the human immunodeficiency virus type 1 (HIV-1) structural precursor Gag protein interacts with the miRNA processing enzyme Dicer. RNA immunoprecipitation and sequencing experiments show that Gag modifies the retention of a specific miRNA subset without affecting Dicer’s pre-miRNA processing activity. Among the retained miRNAs, miR-642a-3p shows an enhanced occupancy on Dicer in the presence of Gag and is predicted to target AFF4 mRNA, which encodes an essential scaffold protein for HIV-1 transcriptional elongation. miR-642a-3p gain- or loss-of-function negatively or positively regulates AFF4 protein expression at mRNA and protein levels with concomitant modulations of HIV-1 production, consistent with an antiviral activity. By sequestering miR-642a-3p with Dicer, Gag enhances AFF4 expression and HIV- 1 production without affecting miR-642a-3p levels. These results identify miR-642a-3p as a strong suppressor of HIV-1 replication and uncover a novel mechanism by which a viral structural protein directly disrupts an miRNA function for the benefit of its own replication.

**IMPORTANCE:** Virus-host relationships occur at different levels and the human immunodeficiency virus type 1 (HIV-1) can modify the expression of microRNAs in different cells. Here, we identify a virus-host interaction between the HIV-1 structural protein Gag and the miRNA-processing enzyme Dicer. Gag does not affect the microRNA processing function of Dicer but affects the functionality of a subset of microRNAs that are enriched on the Dicer-Gag complex compared to on Dicer alone. We show that miR-642a-3p, the most enriched microRNA on the Dicer- Gag complex targets and degrades AFF4 mRNA coding for a protein from the super transcription elongation complex, essential for HIV-1 and cellular transcription. Interestingly, the silencing capacity by miR-642a-3p is hindered by Gag and heightened in its absence, consequently affecting HIV-1 transcription. These findings unveil a new paradigm that a microRNA function rather than its abundance can be affected by a viral protein through its enhanced retention on Dicer.

## INTRODUCTION

Interactions between the RNA interference (RNAi) pathway and human immunodeficiency virus type 1 (HIV-1) have been extensively studied, but the systemic outcomes of these interactions remain poorly understood. RNAi is a critical regulatory system present in most eukaryotes, which shapes the transcriptome for mammalian development and homeostasis as well as disease through micro (mi)RNAs and other short non-coding (nc)RNAs (1–3). During miRNA biogenesis, Dicer, an endoribonuclease associated with TAR RNA-binding protein (TRBP) captures precursor (pre-)miRNAs, removes their terminal loops, cleaves the double-stranded (ds)RNA and forms the mature miRNAs (4–8). Dicer, TRBP and the mature miRNAs associate with one of four mammalian Argonaute proteins (Ago1, Ago2, Ago3 and Ago4) to form RNA-induced silencing complexes (RISCs). Within the RISC, Ago proteins remove the passenger strand of the miRNA so that the intact antisense or guide strand can associate with complementary mRNA sequences. The Ago-guide strand-mRNA complexes then associate with several effector enzymes to form cytoplasmic foci called GW bodies (GWBs), which mediate translational repression and mRNA decay (2, 9–11).

HIV-1 proviral DNA integrated into a host chromosome can remain silent or be transcribed by recruiting RNA polymerase (Pol) II. RNA Pol II is subject to transient post-initiation pausing at the promoter-proximal regions of most human genes (12, 13). Release of RNA Pol II from promoter-proximal pausing at genes including HIV-1 requires the kinase activity of positive transcription elongation factor b (P-TEFb), which is tightly regulated in the nucleus through sequestration by the 7SK small nuclear ribonucleoprotein (7SK snRNP).

P-TEFb activity on the promoter is amplified by associating with other kinases on a dimerized scaffold of AF4/FMR2 Family Members 1 & 4 (AFF1/4) to form the human super elongation complex (SEC)(14–17). As promoter-proximal pausing is a critical barrier to HIV-1 expression, the viral *trans*-activator of transcription (Tat) has evolved to both liberate P-TEFb from the 7SK snRNP and facilitate the assembly of hyperactive SECs that preferentially accumulate at paused nascent HIV-1 RNAs for profound viral amplification (14, 18–23). Therefore, the expression or depletion of members of the SEC, such as AFF4, can also have profound effects on HIV-1 transcription and viral production (17, 24–27).

Proteomics suggest distinct interactions between HIV-1 and numerous cellular proteins including Dicer, TRBP and Ago2 in human cells (28, 29). Indeed, the HIV-1 Gag protein recruits Ago2 to RNA packaging signals on unspliced retroviral RNAs to aid in particle packaging without stimulating miRNA-mediated translational repression of these genomic viral RNAs (30). In contrast, the HIV-1 accessory protein Vpr stimulates the proteasomal degradation of Dicer in macrophages by recruiting the ubiquitin ligase E3 complex to the endoribonuclease, thereby enhancing HIV-1 infection in these cells (31). Specific human miRNAs have also been shown to have either positive or negative effects on HIV-1 expression by directly targeting accessible regions in HIV-1 RNAs, or by targeting cellular factors that prevent or contribute to virus replication (32–35).

To understand how HIV-1 interferes with host regulatory processes that influence viral expression, we explored interactions between viral proteins and cellular components of the RNAi pathway. We show that although RNAi activity remains largely unchanged in HIV-1-expressing cells, Gag interacts with Dicer and modulates its association with a specific subset of miRNAs. Among these miRNAs, miR-642a-3p shows an enhanced occupancy on Dicer in the presence of Gag and is predicted to target AFF4 mRNA. We demonstrate experimentally that miR-642a-3p directly downregulates AFF4 expression leading to the inhibition of HIV-1 production. We further demonstrate that Gag expression in cells modulates miR-642a-3p’s effect on AFF4, thereby increasing HIV-1 expression and amplifying viral replication.

## RESULTS

### RNAi remains functional and RNAi proteins are not re-localized in cells expressing HIV-1

Conflicting evidence suggests that HIV-1 components either suppress RNAi or have no substantial effect on the system in mammalian cells (36–41). To resolve this issue, we first evaluated whether RNAi remains functional in cells expressing HIV-1. We assayed EGFP expression in the presence of a GFP-targeting short hairpin (sh)RNA (shGFP) or an exogenous miRNA (miGFP) targeting the transcript, which we previously have shown to be functional in HeLa cells and sensitive to RNAi dysfunction in TRBP knock-out and knock-down cells (42). We found that HIV-1 expression from proviral DNA, clone pNL4-3, had no substantial effect on the extent of EGFP knockdown by shGFP or miGFP in comparison to a control plasmid (pcDNA3.1) (Fig. 1A). Similarly, using an shRNA targeting GAPDH, we found that HIV-1 expression had no major effect on silencing GAPDH transcripts, whether they were expressed from a transfected plasmid (GAPDH-PL) or from the endogenous gene (GAPDH) (Fig. 1B).

**Figure 1:**
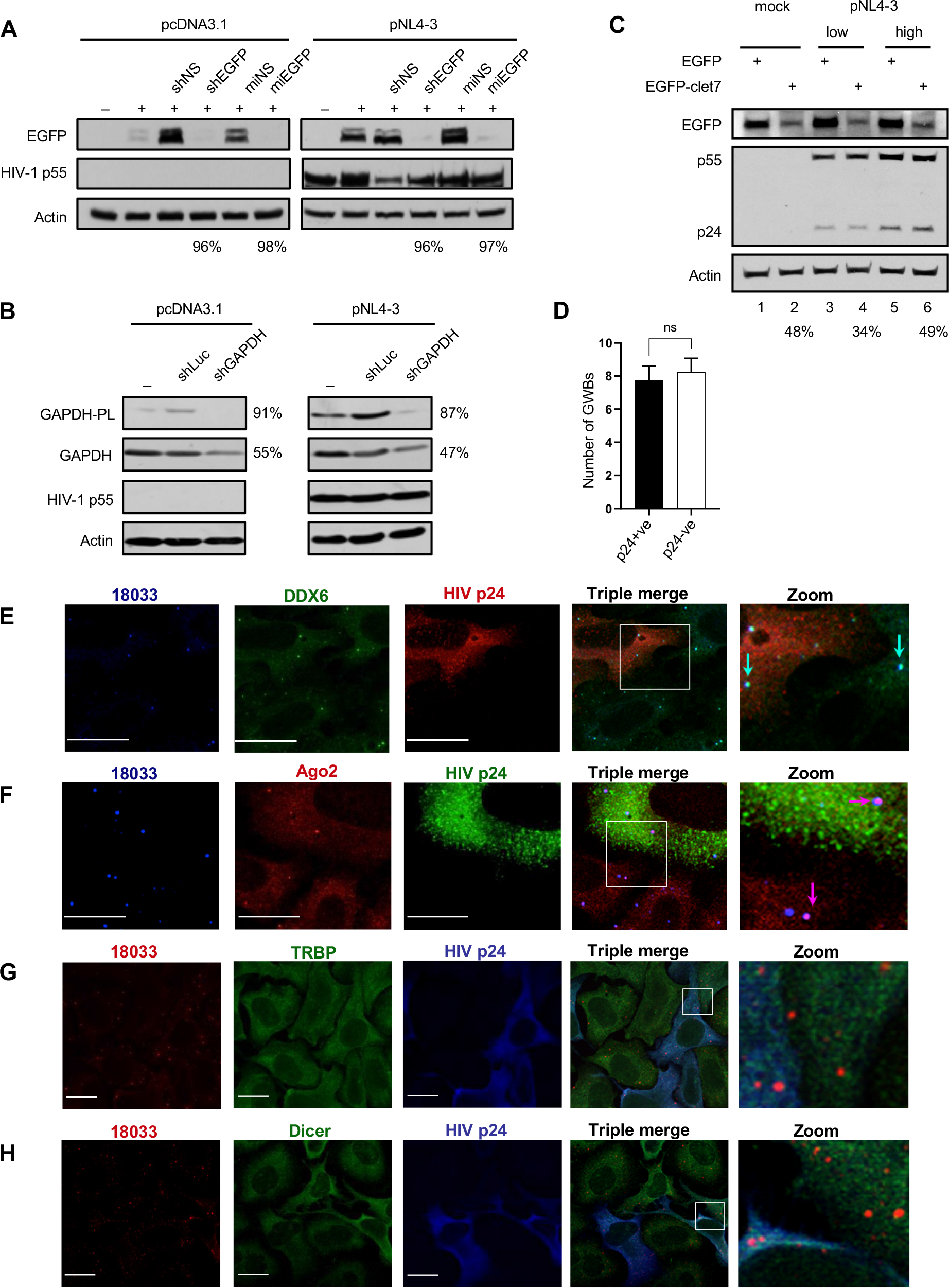
RNAi is functional and the localizations of its contributors remain unchanged in HIV-expressing cells. (A) Silencing of EGFP expression by shGFP and miGFP remains functional in HIV-1 expressing cells. HeLa cells were co- transfected with 2 µg of pcDNA3 or pNL4-3, 1 µg of pEGFP-C1 and 4µg of shRNA-NS, shRNA-EGFP, miRNA-NS or miRNA-EGFP, as indicated. EGFP, HIV-1 Gag and actin expression were observed by immunoblot and EGFP silencing effects in relation to paired nonsense controls were quantified by densitometry analysis. (B) Inhibition of endogenous and transfected GAPDH expression by shGAPDH is functional in HIV- expressing cells. HeLa cells were co-transfected with 2 µg pcDNA3 or pNL4-3, 1 µg GAPDH-ProLabel and 1 µg pSIREN-shRNA-LucIRR or pSIREN-shRNA-GAPDHB. GAPDH, HIV-1 Gag and actin expression were observed by immunoblot and GAPDH silencing effects in relation to paired nonsense controls were quantified by densitometry analysis. (C) Inhibition of EGFP expression by let7 is functional in HIV-1 replicating cells. HeLa-CD4 (MAGI) cells were not infected (lanes 1, 2) or infected with pNL4-3 at one million RT cpms (lanes 3, 4) or 2 million RT cpms (lanes 5, 6) of supernatant. They were co-transfected with pEGFP or pEGFP-clet7 24 h later. EGFP, HIV-1 Gag and actin expression were observed by immunoblot and EGFP silencing effects in relation to paired nonsense controls were quantified by densitometry analysis. (D) The number of large GWBs is not altered in HIV-1 expressing cells. HeLa cells were transfected with 0.2 µg of HIV-1 pNL4-3. Cells were fixed and stained 48 h later with the human GWB marker serum 18033 and an antibody targeting HIV-1 p24. Side-by-side images of p24 positive (+ve) and negative (-ve) cells were identified and images were adjusted to visualize only large 18033 stained GWBs, which were then counted. The bar graph shows the mean number of GWBs from 12 pairs of cells. Statistical analysis was performed using an unpaired Mann-Whitney test (p=0.7) between +ve or –ve cells. **(E-H) The localization of miRNA biogenesis enzymes and RNAi proteins in the cytoplasm is not altered in HIV-1 expressing cells**. HeLa cells were transfected, fixed and stained as in (D) with the addition of antibodies against DDX6 (E), Ago2 (F), TRBP (G) or Dicer (H). The size scale is 20 µm and is shown in each picture on the bottom left. Digitally zoomed images of the merged channels are shown on the far right, with arrows highlighting colocalization between the GWB marker 18033 and DDX6 (E) or Ago2 (F), shown as cyan and purple dots, respectively.

To assess HIV-1’s effect on RNAi mediated by endogenous miRNAs, we used a construct in which the complementary sequence of a human let-7 miRNA is inserted into the 3’UTR (untranslated region) of EGFP (36). We assessed the knock-down of EGFP by the endogenous miRNA let7 in HeLa-based MAGI cells mock or infected by HIV-1 NL4-3 (Fig. 1C). Again, no difference was observed in the efficiency of EGFP knock-down by let-7, indicating that endogenous miRNA biogenesis and RNAi function remain largely unaffected in the presence of replicating HIV-1.

We next evaluated whether the virus might exert an effect on the cellular localization of proteins involved in RNAi. HeLa cells were transfected with HIV-1 pNL4-3 and incubated for 48 h before being assayed for immunofluorescence (IF) under a confocal microscope. IF assays were performed using antibodies targeting the HIV-1 capsid (p24) and its Gag precursor protein (p55), the human autoimmune serum 18033 as a GWB marker recognizing the enhancer of mRNA decapping 4 (EDC4, formally known as Ge-1) and GW182 (43, 44), combined with either TRBP, Dicer, Ago2 or the GWB-enriched translational repressor DDX6 (9, 45). We did not observe any obvious differences in the size or number of GWBs between side-by-side images of p24 positive and negative cells (Fig. S1) and there were no significant differences in the number of large GWBs (Fig. 1D). We then verified the presence and abundance of RNA silencing effector proteins in HIV- 1-expressing and non-expressing cells and found that both DDX6 and Ago2 colocalized to GWBs in p24 positive and negative cells (Fig. 1E, F). The cytoplasmic localization of TRBP and Dicer with a preferential cytoplasmic distribution outside of GWBs were also similar between p24 positive and negative cells (Fig. 1G, H). We also noticed a partial colocalization between Dicer and Gag, observed as cyan fluorescence, in HIV-1-expressing cells (Fig. 1H, Zoom). We conclude that the cytoplasmic distribution of GWBs, TRBP, Dicer, Ago2 and DDX6 remains unchanged in HIV-1 expressing HeLa cells, but that Gag partially colocalizes with Dicer.

### Viral Gag colocalizes with Dicer in HIV-1 producing cells

To confirm and further evaluate the colocalization between Dicer and HIV-1 Gag observed in Fig. 1H, we performed additional IF analyses over several time points using a different antibody for Dicer. We transfected cells with pNL4-3 and observed a colocalization between Dicer and Gag at 24, 36, 48 and 72 h post-transfection (Fig. S2). The colocalization was most notable at 48 h, with a Pearson’s correlation coefficient of *r* =0.6380 ± 0.0290, indicating a proximity and suggesting a possible interaction between the two proteins (Fig. 2A).

**Figure 2:**
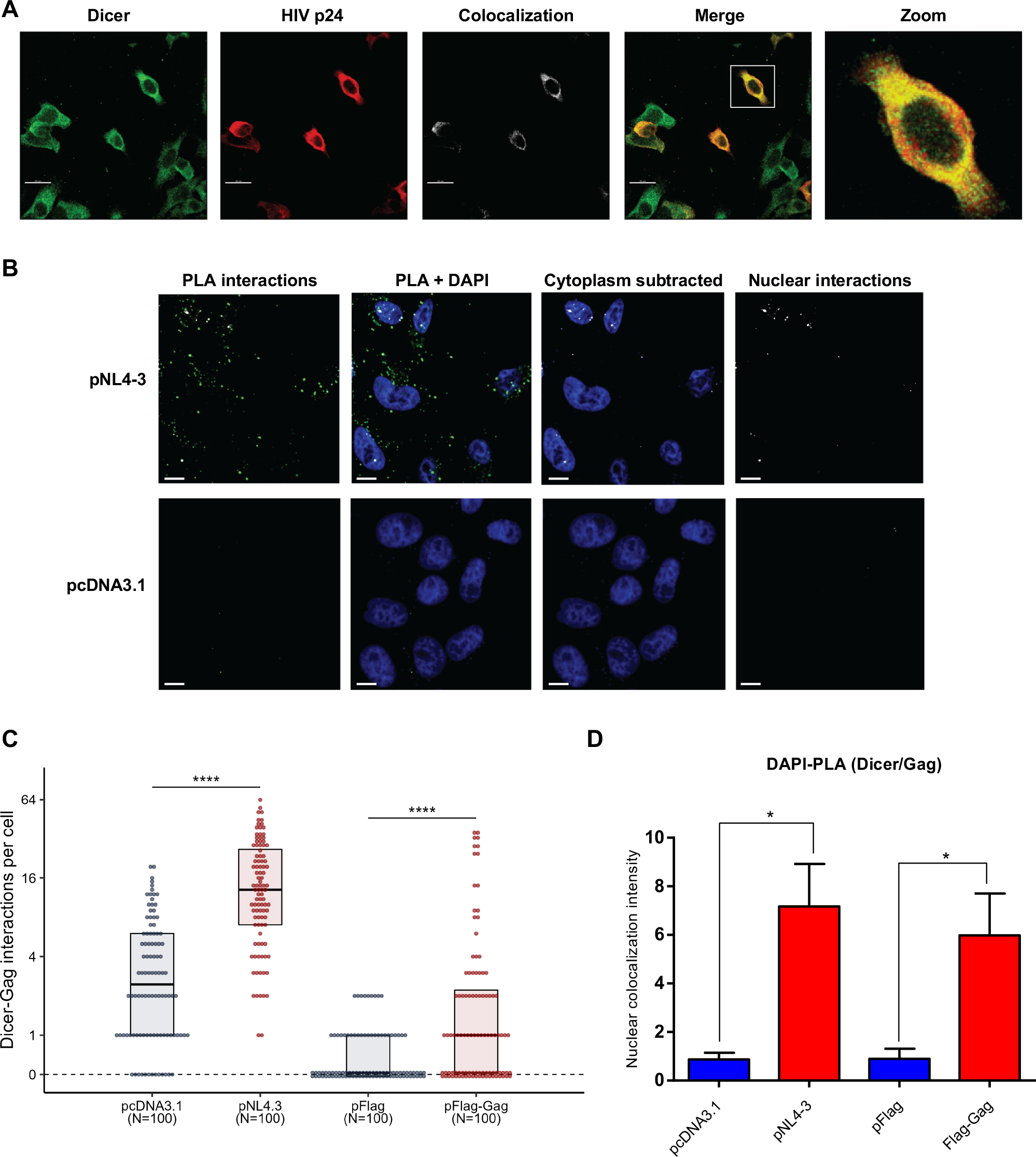
The HIV-1 polyprotein Gag and the endogenous human protein Dicer associate together in cells. (A) Dicer and Gag colocalization. HeLa cells were transfected with 0.2 µg of HIV-1 pNL4-3 and harvested at 48 h before being fixed and stained with mouse anti-Dicer 13D6 (green) and rabbit anti-p24 (red). A 20 µm scale is shown at the bottom left of each image. Digitally zoomed images of the merged channels are shown on the far right. A colocalization channel was built using Imaris software and is displayed in the third lane. The calculated Pearson’s correlation coefficient is the average from 5 cells ± standard error of the mean (SEM) and is: 0. 0.6380 ± 0.0290 at 48 h. **(B) Dicer and Gag PLA.** Images of HeLa cells transfected with pcDNA 3.1 (bottom) or pNL4- 3 (top) are representative of 100 counted cells. A 10 µm scale is shown at the bottom left of each image. From left to right: the first column shows a representative field in the PLA channel. The second column shows the merged channel of PLA and DAPI. The third column shows the merged channel, only including PLA signals within the nucleus. The fourth column shows the colocalization signal between PLA and DAPI (interactions within the nucleus). **(C) PLA boxplot (log2) of Dicer-Gag interaction events.** PLA quantification of Dicer-Gag interactions within HeLa cells. Statistical analysis was performed using an unpaired Mann-Whitney test: pcDNA3.1 versus pNL4-3, p < 0.0001 and pFlag versus Flag-Gag p < 0.0001. **(D) Dicer-Gag colocalization within the nucleus.** Graph comparison of nuclear colocalization intensity of DAPI-PLA (Dicer-Gag) with pcDNA3.1 versus pNL4-3 (n>10, p=0.0273) and pFlag versus Flag-Gag (n>10, p=0.0101). Statistically significant differences between groups were assessed by *t*-test.

We subsequently characterized complex formation between Dicer and Gag by performing *in situ* proximity ligation assays (PLA). Using PLA with p24 and Dicer antibodies and fluorescent probes in pNL4-3-transfected HeLa cells, we observed a substantial number of proximity events in cells containing HIV-1 and few baseline signals in mock-transfected cells (Fig. 2B). We obtained a similar result when we transfected HeLa cells with a Flag- Gag vector (Fig. S3). We quantified the number of PLA interactions in 100 cells for each condition, obtaining up to 64 Dicer-Gag proximity events in individual pNL4-3-transfected cells and up to 36 in Flag-Gag-transfected cells (Fig. 2C). We also evaluated Dicer-Gag colocalization in the nucleus, viewed as white dots within DAPI-stained regions (Fig. 2B, Fig. S3), by removing cytoplasmic dots and quantifying the mean fluorescence intensity in the merged DAPI-PLA channel. Cells containing HIV-1 or Flag-Gag had a higher colocalization intensity than mock-transfected cells (n>10, P< 0.01) (Fig. 2D). We conclude that Gag comes into close proximity with Dicer predominantly in the cytoplasm and to a lesser extent in the nucleus.

### Gag interacts with Dicer without affecting its catalytic activity

To confirm whether Dicer and Gag interact, we transfected human embryonic kidney (HEK) 293T cells with pNL4-3 and performed immunoprecipitations (IP) with a Dicer antibody. Our co-IPs showed that endogenous Dicer pulled down Gag from cells expressing HIV-1, which was not observed with an isotype antibody (Fig. 3A). To further determine the specificity of this interaction, we co-transfected HEK 293T with GST-Dicer and either pFlag-Gag or the empty vector pFlag. Co-IP with a GST antibody showed a 55 kDa band corresponding to Flag-Gag (Fig. 3B), confirming that Gag forms a complex with Dicer. Treatment with RNases before GST co-IP did not prevent pulling down Gag, showing that the Dicer-Gag interaction does not depend on an RNA mediator (Fig. 3C). Dicer’s interaction partners TRBP and PACT have previously been shown to differentially affect substrate RNA selection, RNA cleavage activity and Ago strand loading (7, 46). Wehypothesized that in interacting with Dicer, Gag might also affect its endoribonuclease’s activity. We thus assessed Dicer’s nuclease activity by incubating immunoprecipitated GST-Dicer with α-^32^P-labelled uridine triphosphate (UTP)-labeled pre-miRNA-let7c in physiological conditions, as previously described (47) (Fig. 3D). In addition, we synthesized pre-miR-29a, which is a regulator of HIV-1 expression and latency (48).

**Figure 3:**
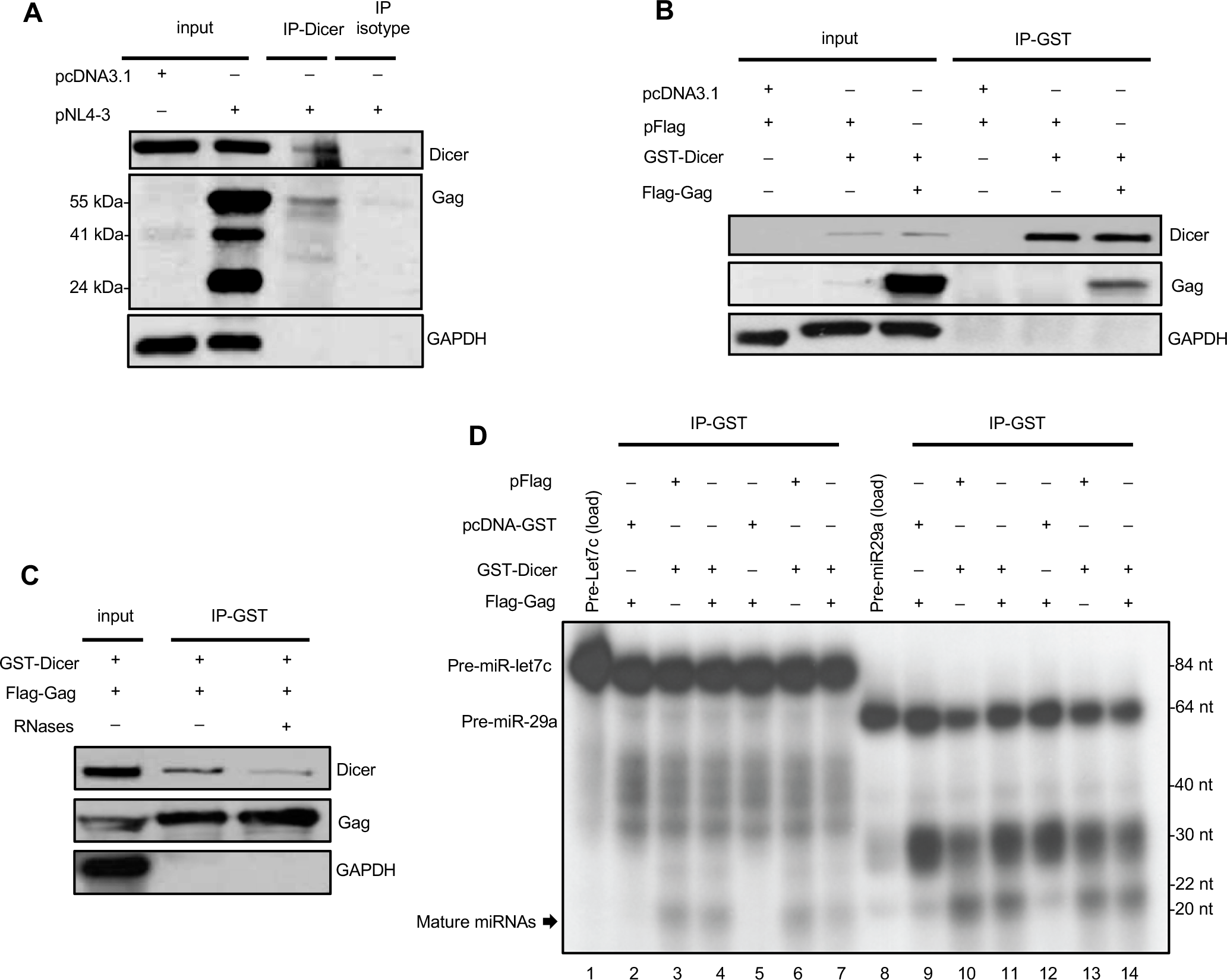
Gag and Dicer interact independently of RNA binding partners and this interaction does not impair Dicer cleavage activity. (A) Endogenous Dicer IP during HIV-1 replication. HEK 293T cells transfected with either pNL4-3 or pcDNA 3.1 for 48h were subjected to IP-Dicer. Gag and Dicer are observed in the second lane (input) as well as in the third lane (output), although Gag is not immunoprecipitated by the isotype control (fourth lane). GAPDH is shown as a loading and IP control. **(B) IP-Dicer during Gag expression**. HEK 293T cells were co-transfected for 48h with mock Dicer (pcDNA3.1)/pFlag, GST-Dicer/pFlag or GST-Dicer/Flag-Gag. The third lane shows the co-expression of Dicer and Gag in the input. Lane 6 shows the interaction between Dicer and the Gag in an IP-GST. **(C) The Dicer-Gag interaction is RNA-independent**. HEK 293T cells were co-transfected with GST-Dicer/Flag-Gag and RNases were added during the IP (lane 4). **(D) Dicer and Dicer-Gag catalytic activity on pre-miR-let7c and pre-miR-29a**. Radiolabeled pre-miRNAs let-7c (lanes 1-7) and 29a (lanes 8-14) were incubated with GST-Dicer (lanes 3,4,6,7, 10,11, 13, 14) and Flag-Gag (lanes 2, 4, 5, 7, 9, 11, 12, 14) and an IP-GST was performed and evaluated for RNA cleavage. Lanes 5-7 are duplicates of lanes 2-4. Lanes 12-14 are duplicates of lanes 9-11. The arrow on the left shows the expected 20-22 nt Dicer-cleaved miRNAs (lanes 3, 4, 6, 7, 10, 11, 13, 14).

Labeled pre-miR-let7c (Fig. 3D, lanes 1-7) and pre-miR-29a (lanes 8-14) were used to assess Dicer’s cleavage of both pre-miRNAs. Incubation of either pre-miRNA with GST- Dicer gave rise to similar levels of 22-nucleotide (nt) mature miRNAs in both the absence (lanes 3, 6, 10, 13) and presence (lanes 4, 7, 11, 14) of Flag-Gag. The 35-50-nt and 30- nt intermediate bands for miR-let7c and miR-29a respectively are likely spontaneous cleavage products not generated by Dicer. These results suggest that the production of mature miR-let7c and miR-29a by Dicer is maintained in the presence of Gag.

### Gag increases the occupancy of specific miRNAs on Dicer

Although HIV-1 does not cause any major disruption in miRNA biogenesis or RNAi function for tested RNAs, our results do not preclude the possibility that Gag affects the occupancy (through modulated pre-miRNA selection or Ago loading) of specific miRNAs on Dicer. To test this hypothesis, we performed IPs with a Dicer antibody or an isotype control from cells expressing Dicer (HA-Dicer) and/or Gag (Flag-Gag) and collected immunoprecipitated RNAs for sequencing (RIP-seq) (Fig. 4A). To identify the unique sets of RNAs that were associated with Dicer in each condition, sample peaks were firstly called using MACSv2 (49) producing 787 peaksets for Dicer and 937 for Dicer/Gag in pooled analyses. We next determined their potential occupancy in Dicer using Diffbind (50) obtaining consensus peaksets of 175 RNAs in HA-Dicer and 121 RNAs in HA-Dicer/Flag-Gag (Fig. 4B). 70 peaks were found in both conditions, 105 RNAs were specific to HA-Dicer and 51 to HA-Dicer/Flag-Gag (Table S1, S2 and S3).

**Figure 4:**
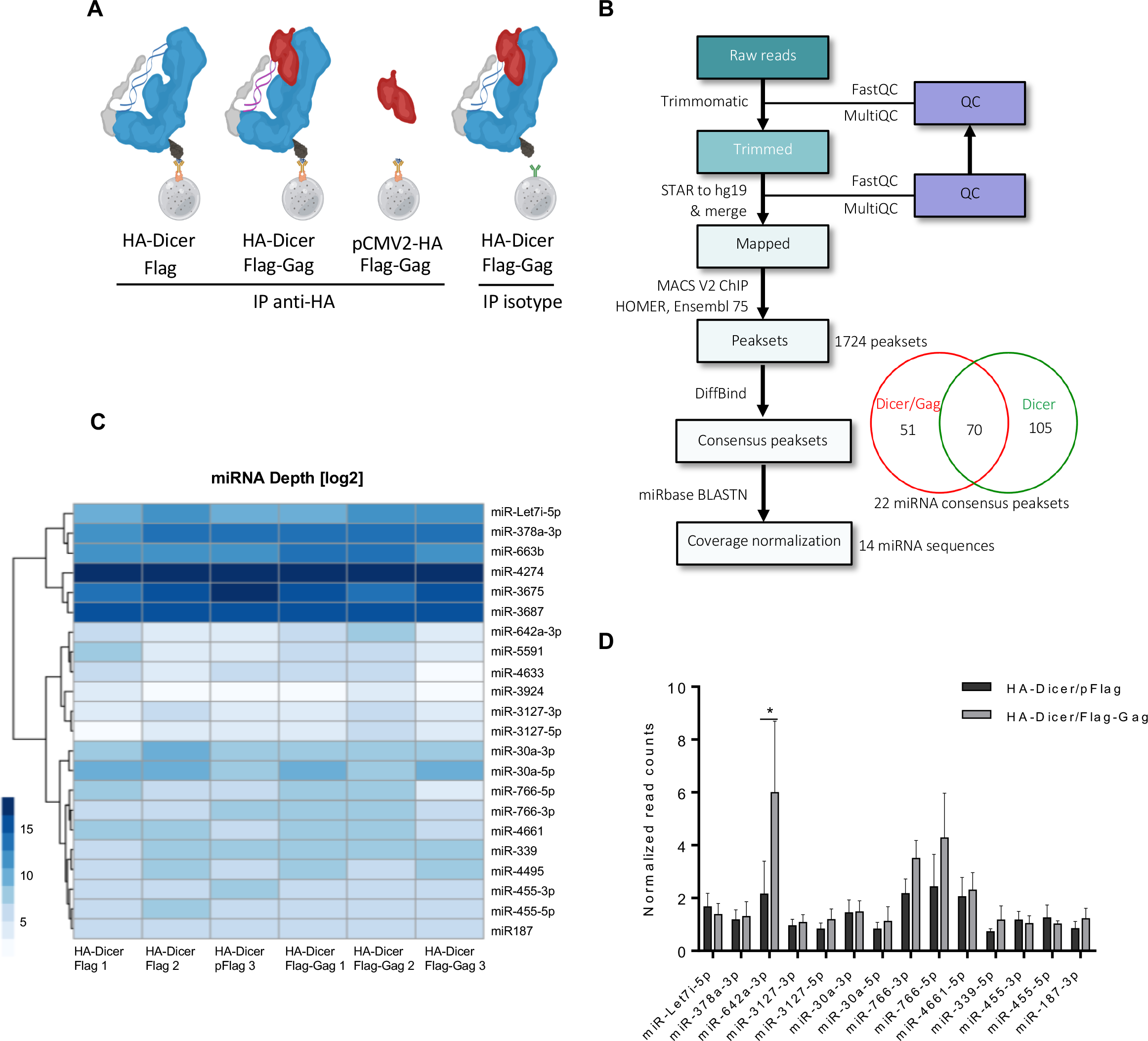
The Dicer-Gag interaction results in an increased occupancy of specific miRNAs on Dicer. (A) Illustrated Dicer RIP-Seq conditions. HEK 293T cells were transfected with1 µg of pCMV2-HA or HA-Dicer, and 1 µg of either pFlag or Flag-Gag for 48 h as indicated. IP assays were carried out for each condition using anti-HA or an isotype antibody as indicated. **(B) Flow chart depicting Dicer and Gag RIP-seq bioinformatic’s pipeline.** Raw reads were trimmed and mapped to produce peaksets. Diffbind was used to determine the occupancy of the peaksets generating sequences annotated as miRNAs. Consensus miRNAs sequences from occupancy analyses were blasted in miRBase and only miRNAs that matched perfectly with IDs and sequences in the database were further normalized to the coverage of controls. **(C) Heat map of sequenced miRNAs.** A log2 heat map of miRNA read depth per replicate in IP-HA- Dicer/pFlag or IP-HA-Dicer/Flag-Gag was generated in R studio. Darker blue colors represent more miRNA reads (see scale). **(D) miRNAs differentially bound to Dicer or Dicer-Gag**. Bar graph of true normalized miRNAs. Data represent the mean ±SEM. A two-way ANOVA with Sidak’s multiple comparison test was carried out for HA-Dicer/pFlag versus HA-Dicer/Flag-Gag with all p values ≥0.05 except for miR-642a-3p (p≤0.05).

Dicer has been shown to also bind and cleave several types of ncRNAs, including transfer RNAs, small nuclear RNAs, small nucleolar RNAs and some long ncRNAs (51), which was consistent with our results. We retrieved 18 ncRNAs annotated as miRNAs (Table S1, S2 and S3). To corroborate the annotation from Diffbind on miRNAs, we aligned our reads to *Homo sapiens* GRCh37 (hg19) using IGV (52–54) and retained only miRNAs from all ncRNAs. Both strands of miR-3127, miR-30a, miR-766 and miR-455 and one strand of the others were present, yielding a total of 22 miRNAs in our final occupancy analysis (Fig. 4C). To ensure the annotated miRNAs corresponded to true miRNAs, we next performed a BLAST search with the 22 miRNA consensus sequences on miRBase (55, 56), resulting in fourteen miRNAs that matched perfectly with sequences and IDs on miRBase.

To determine quantitative differences between the resulting miRNAs in HA-Dicer and HA- Dicer/Flag-Gag, we normalized each miRNA’s coverage of samples and controls to the total number of reads followed by a comparison to the mean coverage for HA/Flag-Gag and the isotype control (Table S4). The differential occupancy showed an overall tendency of more miRNAs bound to Dicer-Gag than to Dicer alone (Fig. 4D). Specifically, miR-642a- 3p, miR-187-3p, miR-30a-5p, miR-766-3p, miR766-5p, miR-3127-5p, miR-339-5p were distinctly more detected on Dicer-Gag, whereas miR-455-5p, 3p and let7i-5p were slightly more associated with Dicer alone, (Fig. 4D). The final products were statistically evaluated using a two-way ANOVA with Sidak correction for multiple comparisons, identifying miR- 642a-3p as the most significant miRNA preferentially associated to Dicer-Gag (p≤0.05).

### MiRNAs that are highly associated with Dicer-Gag complex are predicted to be involved in virus-regulating functions

We further evaluated the best miRNA candidates among this list based on i) their preferential occupancies in the Dicer/Gag complex (miR-642a-3p, miR-766-5p and miR- 766-3p) or ii) their known involvement in the control of HIV-1 replication [miR-378a-3p (57) and miR-30a-5p (58)]. miRNAs were quantified by reverse transcription quantitative polymerase chain reaction (RT-qPCR)(59) on HA-IP samples from cells co-transfected with HA-Dicer and either pFlag or pFlag-Gag. The normalized expression of each tested miRNA was not significantly different (p>0.05) between Flag and Flag-Gag conditions as measured in whole cell lysates (Fig. 5A-E). However, all 5 miRNAs were statistically more abundant on Dicer in Flag-Gag expressing cells than in control cells (Fig. 5F-J). Specifically, miR-642a-3p showed a significant (p<0.001) three-fold increased occupancy on Dicer in the presence of Gag (Fig. 5F). miR-766-5p and miR-766-3p similarly showed a two-fold increased binding to Dicer in the presence of Gag, which corroborates the trends seen in RIP-Seq data (Fig. 5G, H). Although miR-30a-5p and miR-378a-3p were also more abundant in the Dicer IP when Gag was present (Fig. 5I, J), their relative normalized expression in corresponding inputs were higher in Gag expressing cells (Fig. 5D, E), suggesting that the noted difference in Dicer binding may reflect a difference in expression. In summary, miR-642a-3p, miR-766-5p and miR-766-3p were found to be significantly more abundant on Gag-bound Dicer than they were on Dicer alone, and this differential occupancy could not be attributed to differential expression in Gag expressing cells.

**Figure 5:**
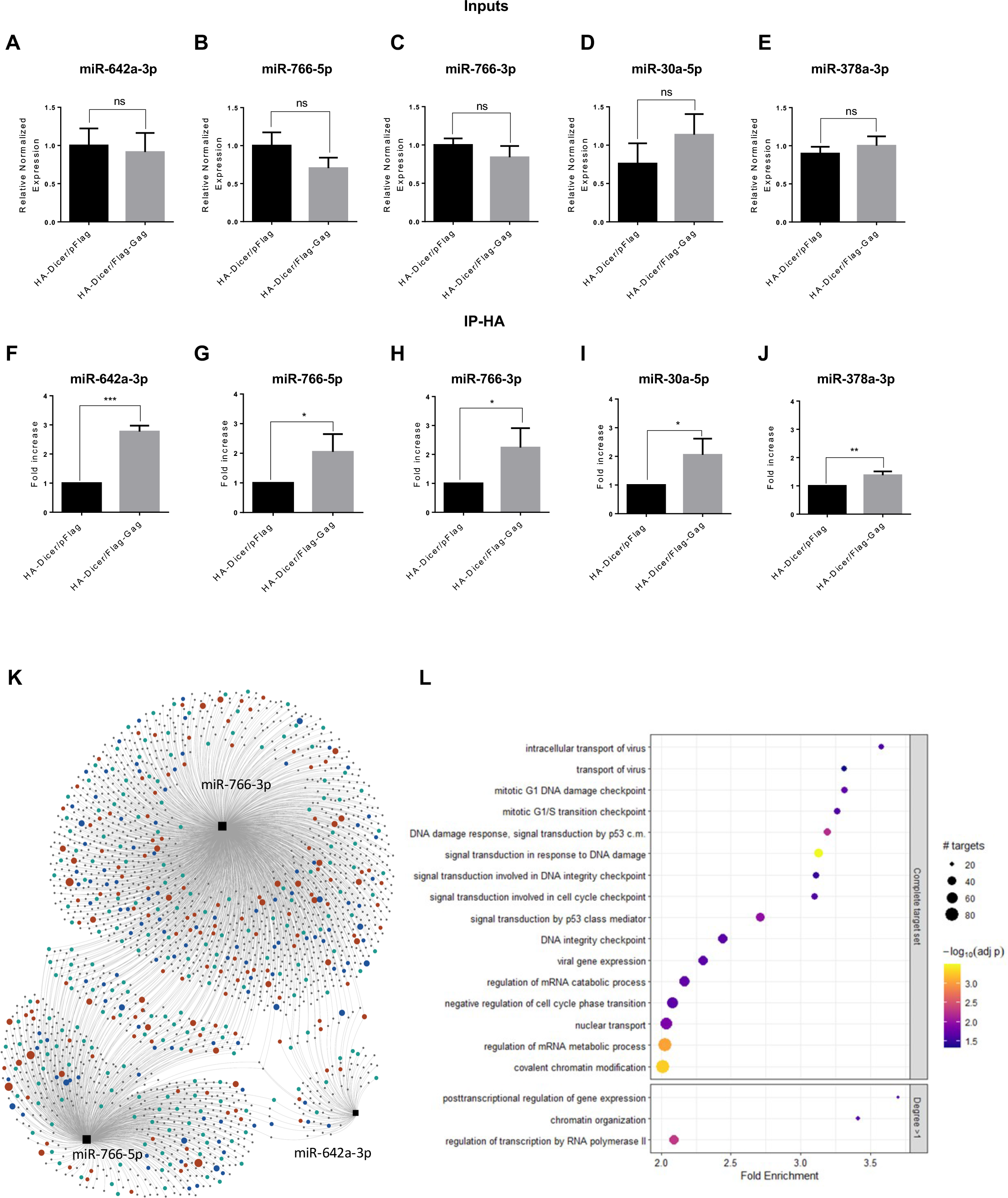
Validated miR-642a-3p, miR-766-5p and miR-766-3p target genes involved in virus-regulating functions. (A-J) RIP-RT-qPCR validation of miRNAs differentially bound to Dicer. The RIP-qRT-PCR validation of miR-642a-3p, miR-766-5p miR-766-3p, miR-30a-5p and miR-378a-3p was carried out in IP-HA-Dicer/pFlag or IP-HA-Dicer/Flag- Gag. miRNA relative normalized expression of the input for (A) miR-642a-3p, (B) miR- 766-5p, (C) miR-766-3p, (D) miR-30a-5p, (E) miR-378a-3p. From (F-J), fold increase of the output (F) miR-642a-3p (p=0.0001), (G) miR-766-5p (p=0.0392), (H) miR-766-3p (p=0.0323), (I) miR-30a-5p (p=0.0254), (J) miR-378a-3p (p=0.0064). Bars represent the mean ±SEM of three independent experiments and *t-*tests were performed to assess significance. **(K) Experimentally curated miRNA-gene interactions on miRTarBase** v.8.0 and TarBase v.8.0. are displayed in a network and annotated using MirNet 2.0. Circles represent genes, squares represent miRNAs and edges represent curated verified interactions. Red nodes represent genes associated with viral processes, blue nodes are involved in the regulation of cell cycle checkpoints, cyan/green nodes are involved in gene expression regulation. Nodes with multiple annotated functions appear larger. **(L) Fisher’s exact test for GO enriched terms**. Enriched GO biological processes annotations for experimentally validated target genes of miR-642a-3p, miR-766-5p and miR-766-3p are sorted by fold enrichment > 2.00 (Fisher’s exact test with Bonferroni correction).

No previous report has suggested that miR-642a-3p, miR-766-5p or miR-766-3p could target HIV-1 RNAs directly. Furthermore, they do not have any predicted targets on the HIV-1 HXB2 reference genome using miRDB’s MirTarget prediction algorithm (60). We therefore explored the endogenous cellular gene targets of the three miRNAs and their relationships to processes known to affect HIV-1 replication. We queried miRTarBase v.8.0 and TarBase v.8.0 (61, 62), two independently curated databases of experimentally identified miRNA-RNA target pairs, to identify targets of the enriched miRNAs. Our results show that miR-766-3p has the highest number of targets and that miR-642a-3p targets a limited number of genes. They also indicate that 30.7% of the genes targeted by miR-642a-3p are shared targets with one or both strand(s) of miR-766-5p/3p, suggesting that these miRNAs have similar targetomes (Fig. 5K). We next investigated biological process gene ontologies (GO) that were enriched among this non-redundant set of target genes against a background of all human miRNA targets listed in miRTarBase and TarBase. Among target biological processes, *intracellular transport of virus* (GO:00075733) was the most enriched term (fold enrichment = 3.58, adjusted p value = 0.0265) (Fig. 5L). Another enriched term, *viral gene expression* (GO:0019080) (fold enrichment = 2.3, adjusted p value = 0.0216) retains little overlap with *intracellular transport of virus*, suggesting that host dependency factors (HDFs) known to be involved in independent stages of viral replication cycles in humans are conserved targets of these three miRNAs. Other enriched functions of the target gene set included multiple processes involved in the regulation of cellular replication by cell cycle checkpoints, as well as gene expression regulation during epigenetic remodeling, transcription or translation pathways (Fig. 5L). We performed the same enrichment analysis on genes in the network that were shared targets of more than one miRNA (degree >1), resulting in three GO terms with more than two-fold enrichment, which were all associated with distinct events in gene expression regulation (Fig. 5L). Overall, Dicer-Gag-enriched miRNAs were predicted to regulate host processes governing events in virus replication, gene expression status and cell replication fate.

To evaluate the functional effects mediated by these miRNAs on HIV-1 replication, we next sought to identify a miRNA-driven virus regulatory pathway that could be characterized experimentally and could represent the general downstream effects of the Dicer-Gag interaction. We ranked all putative human gene targets of the three Dicer-Gag- enriched miRNAs using the online prediction database miRDB (60). The database listed 564 targets for miR-642a-3p, 1043 for miR-766-5p and 916 for miR-766-3p. We tabulated genes whose miRDB targeting scores were above 95/100, including 13 targets for miR- 642a-3p, 21 for miR-766-5p and 15 for miR-766-3p (Tables 1-3). Among these, several host genes encoding proteins that have a known interaction with HIV-1 components (called HIV-1 interactome) in the NCBI HIV-1 human interaction database were present (63–65). We also performed a literature search to determine whether any of the listed genes belong to the HIV-1 interactome or had any known function associated with regulating HIV-1 replication. Among the high-ranking presumptive targets, three miR- 642a-3p targets (Table 1), six miR-766-5p targets (Table 2) and two miR-766-3p targets (Table 3) were associated with HIV-1 replication or the HIV-1 interactome. Notably AFF4 (previously called MCEF), a key scaffold component of the human SEC and known regulator of HIV-1 expression (14, 17, 24, 25, 66) was among the top rated putative targets of miR-642a-3p. AFF4 is also listed in miRTarBase v.8.0 as a target of miR-642a-3p from data collected in two independent high-throughput studies, suggesting that the predicted interaction was reproducible (61, 67). Based on these findings, we further investigated the potential activity of miR-642a-3p on AFF4 mRNA and protein as a downstream effect of the Dicer-Gag interaction.

**Table 1.** High ranked target genes for miR-642a-3p

| Target Rank | Target score | Gene Symbol | HIV-1 interactome | Associated with HIV-1 replication | References |
| --- | --- | --- | --- | --- | --- |
| 1 | 99 | AGO4 |  |  |  |
| 2 | 99 | ZNF449 |  |  |  |
| 3 | 98 | AFF4 | ✓ |  | (16, 70, 108) |
| 4 | 98 | NCOA7 |  |  |  |
| 5 | 98 | DIP2C |  |  |  |
| 6 | 97 | PABPN1 |  | ✓ | (109) |
| 7 | 97 | TMEM116 |  |  |  |
| 8 | 96 | VGLL3 |  |  |  |
| 9 | 96 | PCDH18 |  |  |  |
| 10 | 95 | SPESP1 |  |  |  |
| 11 | 95 | BCL2L2-PABPN1 |  |  |  |
| 12 | 95 | SOD2 |  | ✓ | (110, 111) |
| 13 | 95 | ARSB |  |  |  |

**Table 2.**
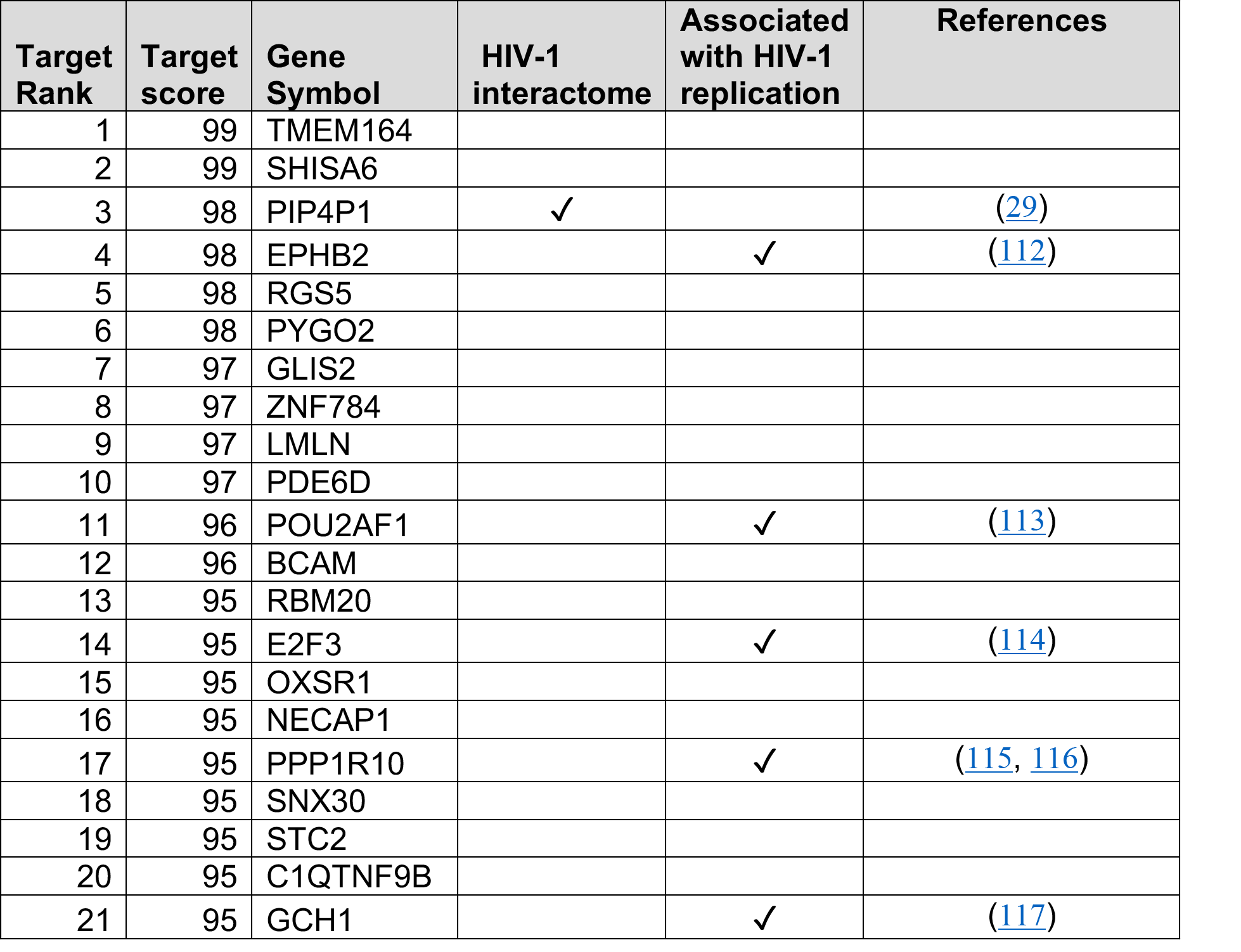
High ranked target genes for miR-766-5p

**Table 3.**
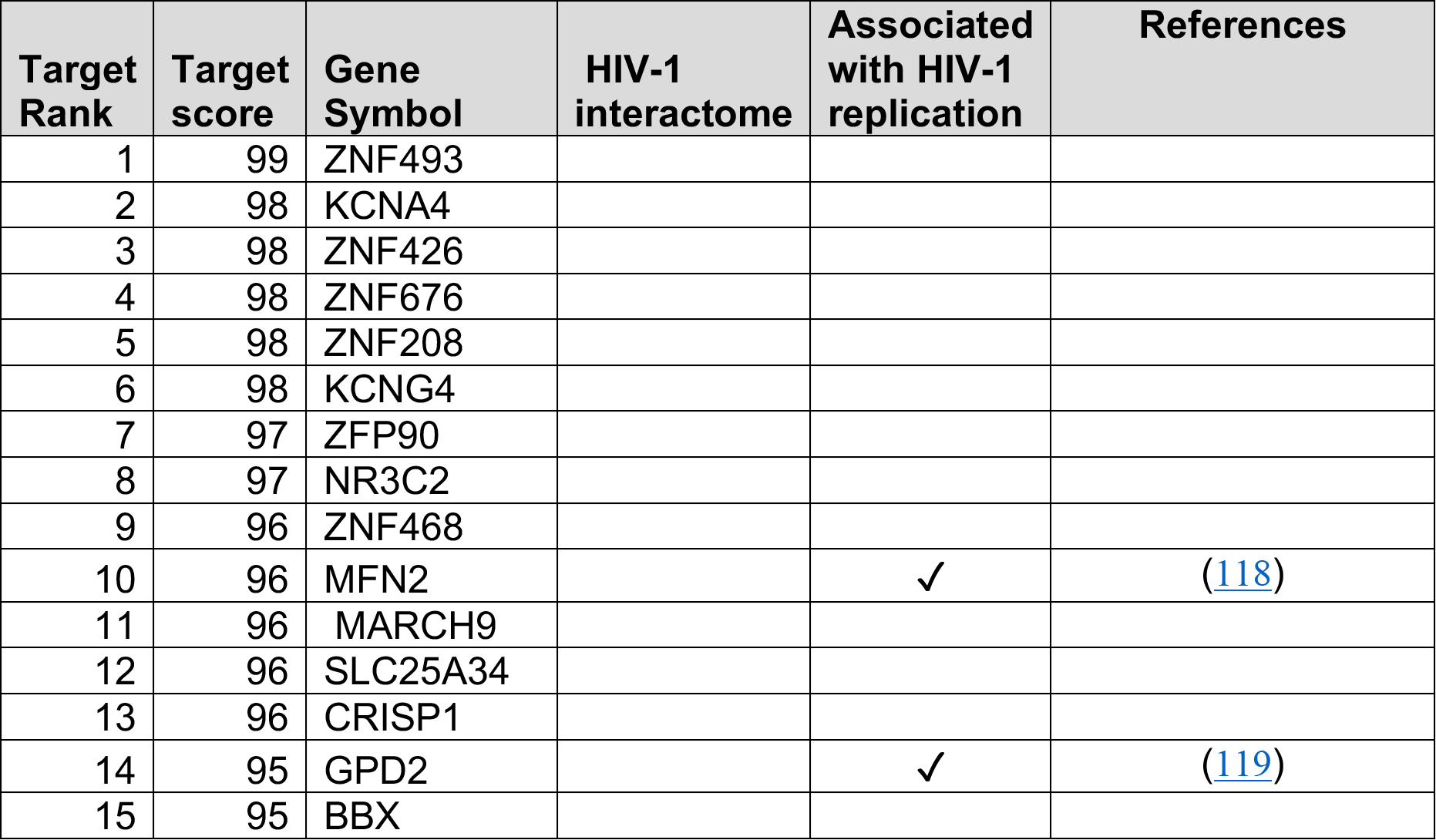
High ranked target genes for miR-766-3p

| Target Rank | Target score | Gene Symbol | HIV-1 interactome | Associated with HIV-1 replication | References |
| --- | --- | --- | --- | --- | --- |
| 1 | 99 | ZNF493 |  |  |  |
| 2 | 98 | KCNA4 |  |  |  |
| 3 | 98 | ZNF426 |  |  |  |
| 4 | 98 | ZNF676 |  |  |  |
| 5 | 98 | ZNF208 |  |  |  |
| 6 | 98 | KCNG4 |  |  |  |
| 7 | 97 | ZFP90 |  |  |  |
| 8 | 97 | NR3C2 |  |  |  |
| 9 | 96 | ZNF468 |  |  |  |
| 10 | 96 | MFN2 |  | ✓ | (118) |
| 11 | 96 | MARCH9 |  |  |  |
| 12 | 96 | SLC25A34 |  |  |  |
| 13 | 96 | CRISP1 |  |  |  |
| 14 | 95 | GPD2 |  | ✓ | (119) |
| 15 | 95 | BBX |  |  |  |

### miR-642a-3p downregulates AFF4 mRNA and protein expression

Before examining how the Gag-mediated selective modulation of miRNA-Dicer occupancy affects AFF4 expression and the HIV-1 replication cycle, we first assessed whether miR- 642a-3p indeed regulates AFF4 mRNA expression. We used TargetScan release 8.0 to output a context score of predicted miRNA interactions at the AFF4 3’UTR according to the known heuristics of human miRNA binding (68, 69). TargetScan is based on a stepwise weighted regression model of known features that regulate miRNA-mRNA target pairing in humans, while miRDB uses a different approach founded upon an independent set of experimental data fed into a support vector machine with no conservation input (60, 68).

miR-642a-3p had the second most favorable cumulative weighted context score for regulation of AFF4 mRNAs of all miRNAs listed on TargetScan, corroborating the interaction identified on miRDB (Table S5).

According to TargetScan and miRDB, AFF4 has eight predicted 7mer miR-642a-3p target sites and was therefore a very likely target for this miRNA. We subcloned TargetScan’s three highest rated sites and their adjacent upstream supplementary pairing sequences from the AFF4 3’UTR into the 3’UTR of EGFP in a pEGFP-C1 vector (Fig. 6A). Compared to a nonsense (NS) miRNA mimic, a miR-642a-3p mimic inhibited EGFP expression from the co-transfected cloned EGFP 3’UTR (Fig. 6B), showing that the predicted sites in AFF4’s 3’UTR are indeed targets of miR-642a-3p. Conversely, an antagomiR of miR- 642a-3p stimulated an increase in EGFP expression compared to a NS antagomiR, showing that there is some endogenous activity of miR-642a-3p against the experimental EGFP 3’UTR in our cellular model which is abrogated by the corresponding antagomiR (Fig. 6B).

**Figure 6:**
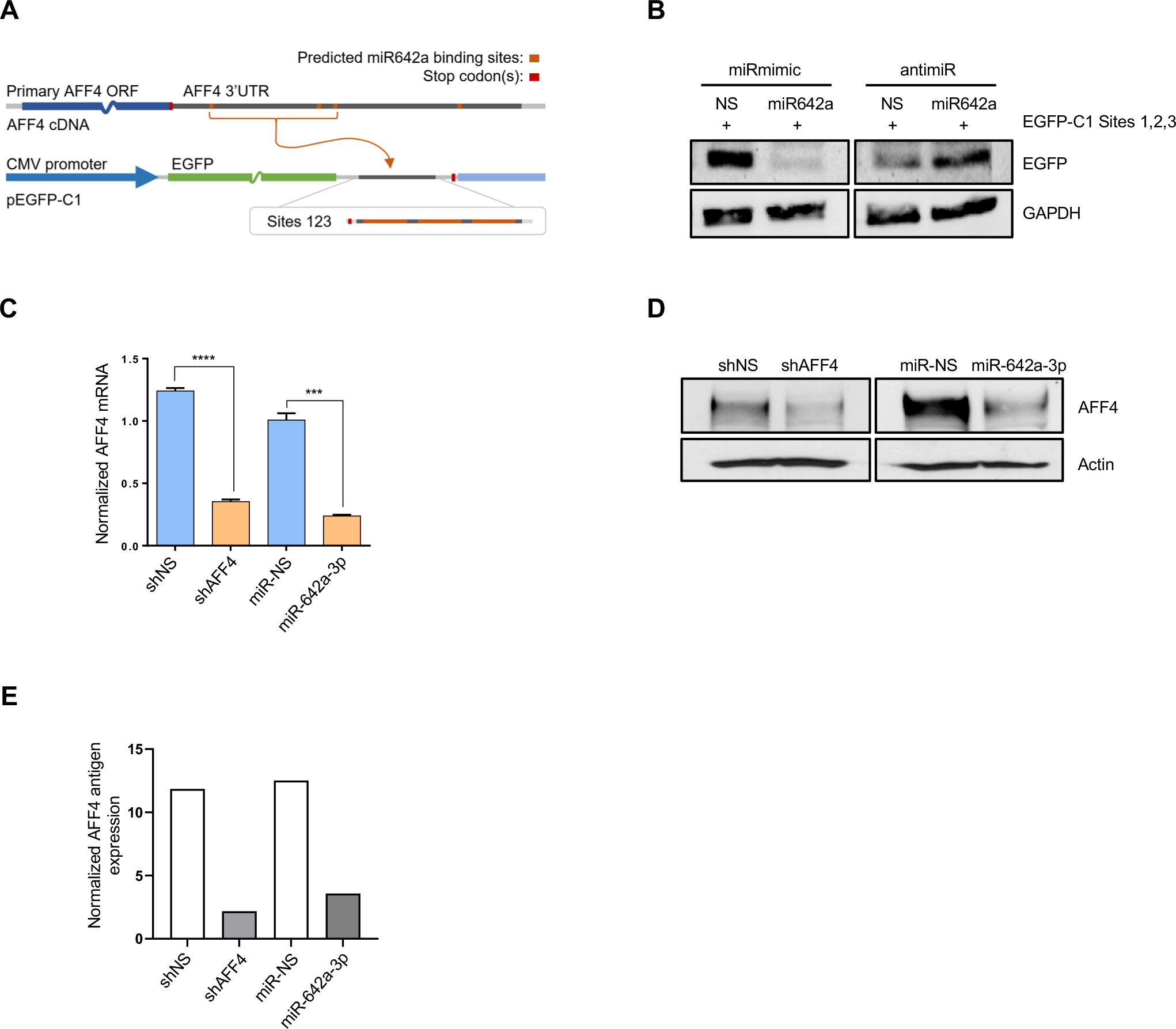
miR-642a-3p modulates AFF4 expression (A) AFF4 predicted binding sites for miR-642a-3p. Schematic showing the location of three TargetScan predicted miR- 642a target sites in the 3’UTR of AFF4 cDNAs that were cloned into the 3’UTR of EGFP in pEGFP-C1 after an inserted Stop codon. **(B) EGFP-C1 is regulated by miR-642a-3p.** HEK 293T cells were transfected with miR-642a-3p or NS miRNA mimics and miR-642a- 3p or NS antagomiRs for 48 h followed by 0.25 μg pEGFP-C1 for 24 h. GAPDH is shown as a housekeeping control. The blot is representative of three independent experiments. **(C) miR-642a-3p silences AFF4 mRNAs.** HEK 293T cells were transfected with shNS, shAFF4 (RNAi positive control), miR-NS or miR-642a-3p, and AFF4 mRNA expression was quantified by RT-qPCR. AFF4 mRNA levels were normalized to actin as an internal control. The graph represents the averages of AFF4 mRNA expression ±SEM of three independent experiments. *t-*tests were performed to assess significance. **(D) miR-642a- 3p silences AFF4 protein expression.** HEK 293T cells were transfected with shNS, shAFF4, miR-NS or miR-642a-3p and AFF4 protein expression was assessed using anti- AFF4-(7D) in a Western blot. GAPDH or actin are shown as a housekeeping controls. The blot is representative of three independent experiments. **(E) Quantification of AFF4 protein expression under silencing conditions.** Bar graph displaying the AFF4 antigen expression in shNS, shAFF4, miR-NS or miR-642a-3p conditions normalized to actin.

We next evaluated the silencing activity of miR-642a-3p on endogenous AFF4 using an shRNA (shAFF4) as a positive control to interfere with mRNA stability or translation of AFF4. HEK 293T cells transfected with shNS or shAFF4 expression plasmids were compared with cells transfected with miRNA mimics miR-NS or miR-642a-3p. After 48 h, we measured AFF4 mRNA expression by RT-qPCR and found that shAFF4 and miR- 642a-3p overexpression significantly reduced AFF4 mRNA abundance in cells by 3.5 fold and 4.2 fold respectively (Fig. 6C). To examine this inhibition at the protein level, a home-made polyclonal antibody raised against amino acids 1-715 was used to recognize both the cloned HA-AFF4 (70) and the endogenous AFF4 protein (Fig. S4). Similarly to the knockdown seen for AFF4 mRNAs, shAFF4 and the miR-642a-3p mimic downregulated endogenous AFF4 protein levels when compared to their individual NS controls (Fig. 6D- E).

### miR-642a-3p downregulates HIV-1 transcription through AFF4, an effect counteracted by Gag

Numerous reports have emphasized the importance of AFF4 in HIV-1 transcription elongation and it has repeatedly been shown that the dysfunction or lack of this protein hinders HIV-1 expression (17, 24–26, 70). Therefore, we hypothesized that knocking down AFF4 by overexpressing miR-642a-3p should reduce HIV-1 expression. To test this hypothesis, we co-transfected HEK 293T cells with HIV-1 pNL4-3 and either miR-NS or miR-642a-3p before quantifying AFF4 mRNAs and elongated HIV-1 transcripts. AFF4 mRNA and HIV-1 transcript levels were respectively reduced by 32% and 33% in the presence of miR-642a-3p (Fig. 7A). HIV-1 protein expression was also evaluated using an antibody targeting the HIV-1 capsid (p24) that also recognizes its precursor Gag in the presence of shAFF4, a miR-642a-3p mimic or a miR-642a-3p antagomiR. shAFF4 and miR-642a-3p reduced, whereas the antagonist of miR-642a-3p slightly increased, HIV-1 expression compared to their respective NS controls (Fig. 7B, C). To better evaluate the antiviral capacity of miR-642a-3p, we compared miR-642a-3p with two miRNAs that are known to downregulate HIV-1 expression: miR-29a-3p and miR-155-5p (48, 71–75). We co-transfected HIV-1 pNL4-3 with equal amounts of miRNA mimics miR-NS, miR-29a-3p, miR-155-5p and miR-642a-3p and found that miR-642a-3p demonstrated slightly more robust HIV-1 silencing activities than did miR-29a and miR-155 (Fig. 7D, E). These miRNAs had no significant impact on cell viability measured by a WST-1 metabolism assay (Fig. 7F).

**Figure 7:**
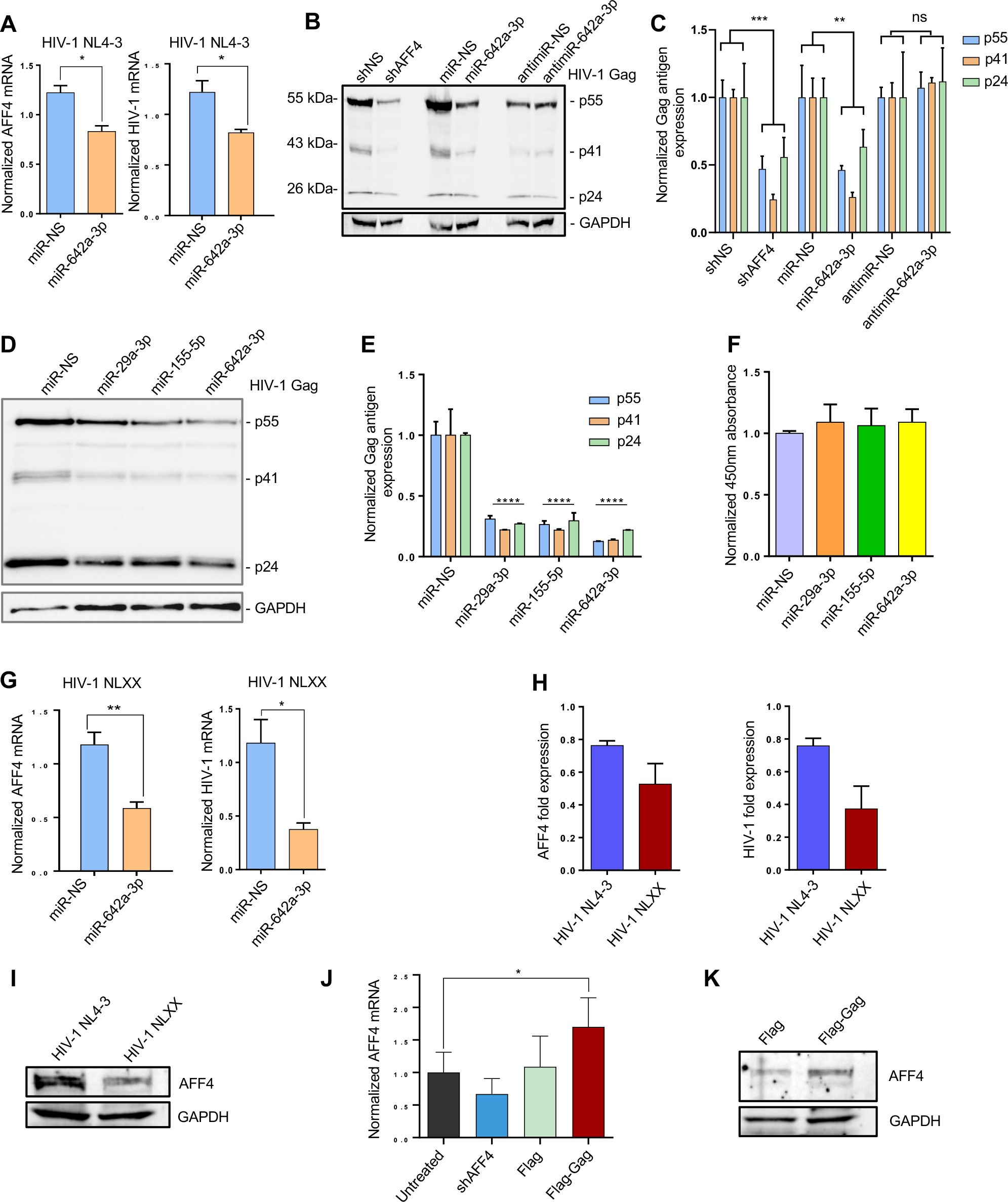
HIV-1 Gag counteracts AFF4 downregulation by miR-642a-3p. (A) miR- 642a-3p downregulates AFF4 and HIV-1 RNA expression. AFF4 (p=0.0134) and HIV- 1 (p=0.0260) mRNA quantification in HEK 293T cells co-transfected with pNL-4-3 and either miR-642a-3p or miR-NS. mRNA was quantified by RT-qPCR and normalized to actin as an internal control. Data are represented as means ±SEM from three independent experiments. *t-*tests were performed to assess significance. **(B) Modulation of HIV-1 protein expression by shAFF4 and miR-642a-3p.** HEK 293T cells were co-transfected with either psiRNA-U6-(shNS or shAFF4) or miRNA mimics (NS or miR-642a-3p) or antimiRs (NS or miR-642a-3p), and 0.5 μg pNL4-3. Whole-cell extracts were subjected to Western blot analysis using anti-p24 and anti-GAPDH antibodies. The blot is a representative of three independent experiments. **(C) Normalized HIV-1 p55, p41 and p24 protein expression in the presence of silencing ncRNA for AFF4.** Expression data was quantified from of Fig 7B and normalized to GAPDH. Data represent the mean ±SEM from three independent experiments. A two-way ANOVA with Tukey’s multiple comparison test (p55, p41, p24): shNS vs shAFF4 (p=0.0008), miR-NS versus miR-642a- 3p (p= 0.0016), antimiR-NS versus antimiR-642a-3p (p= 0.9685). **(D) miRNA mimics miR29a-3p, miR-155-5p, miR-642a-3p decrease HIV-1 expression**. HEK 293T cells were co-transfected with 0.5 μg pNL-4-3 and either miR-NS, miR29a-3p, miR-155-5p or miR-642a-3p. Immunoblot was performed with anti-p24 and anti-GAPDH antibodies. The blot is representative of three independent experiments. **(E) Normalized Gag protein expression in the presence of miRNA mimics**. Expression data quantified from Fig 7D and normalized to GAPDH. Data represent the mean ±SEM from three independent experiments. A two-way ANOVA with Dunnett’s multiple comparison test (p55, p41, p24): miR-NS versus miR-29a-3p (p < 0.0001), miR-155-5p (p < 0.0001), miR-642a-3p (p < 0.0001). **(F) Treatment with miRNA mimics does not affect cell viability.** WST-1 assay on HEK 293T cells treated with miRNAs mimics for 48 h. Data are represented as mean blanked (450 nm/690 nm) absorbances normalized to miR-NS from three independent experiments ±SEM. **(G) miR-642a-3p downregulates AFF4 and HIV-1 (no Gag) mRNA expression.** HEK 293T cells were co-transfected with 0.5 μg HIV-1 pNL-XX (no Gag) and either miR-642a-3p or miR-NS. AFF4 (p=0.0096) and HIV-1 pNL-XX (p=0.0243) mRNA were quantified by RT-qPCR and normalized to actin as an internal control. Data are represented as means ±SEM from three independent experiments. *t-*tests were performed to assess significance. **(H) miR-642a-3p-mediated expression in AFF4 and HIV-1 in the presence or absence of HIV-1 Gag.** Fold expression of AFF4 and HIV-1 following miRNA mimic treatment and transfection with either pNL4-3 (Fig 7A) or pNL-XX (Fig 7F), calculated based on RT-qPCR fold changes in between miR-NS and miR-642a-3p treatments. **(I) AFF4 protein expression during HIV-1 or HIV-1 (no Gag) replication.** HEK 293T cells were co-transfected with 1 μg/mL pNL4-3 or pNL-XX (no Gag) for 48 h. Whole-cell extracts were subjected to Western blot using anti-AFF4 (7D) and anti-GAPDH antibodies. Shown is a representative blot among three replicates. **(J) Gag is sufficient to increase endogenous AFF4 mRNA expression.** RT-qPCR of AFF4 mRNAs normalized to actin, from HEK 293T cells transfected with psiRNAU6-shAFF4, pFlag or Flag-Gag for 48 h. Data are represented as means ± SEM from five independent experiments. An ANOVA (p=0.0046) and a subsequent Dunnett’s multiple comparisons test p(untransfected/Flag-Gag)=0.0262, p(Flag/Flag-Gag)=0.0685 (ns). **(K) Gag expression is sufficient to increase endogenous AFF4 translation.** HEK 293T cells were co-transfected with 1 μg/mL pFlag or Flag-Gag for 48 h. Whole-cell extracts were subjected to Western blot using anti-AFF4 (7D) and anti-GAPDH antibodies. Shown is a representative blot among three replicates.

We finally sought to determine how the enrichment of miR-642a-3p on Gag-bound Dicer might affect the antiviral activity of this miRNA. To address this question, we measured the abundance of AFF4 mRNAs and elongated HIV-1 transcripts in cells transfected by an HIV-1 molecular clone that does not express Gag (pNLXX) (76). In the presence of miR-642a-3p mimics, AFF4 mRNA levels in these cells were reduced by 50% and elongated HIV-1 pNLXX transcripts were reduced by 68% (Fig. 7G). Of note, the endogenous repression of AFF4 observed in the absence of Gag expression from pNLXX (Fig. 7G) was more robust than what was observed when Gag was expressed in the context of pNL4-3 (Fig. 7A). Quantitative comparison showed that the presence of Gag rescued AFF4 mRNA abundance by 33% and elongated HIV-1 transcript abundance by 47% (Fig. 7H). These results suggest that Gag prevents the activity of miR-642a-3p on AFF4 and on downstream HIV-1 transcription. To determine if the observed rescue by Gag could also be observed at the level of AFF4 protein expression under endogenous levels of miR-642a-3p regulation, we transfected cells with HIV-1 pNL4-3 or HIV-1 pNLXX and confirmed that AFF4 was more abundant when Gag was expressed (Fig. 7I). In cells transfected with pFlag or pFlag-Gag, we similarly found that Gag alone was sufficient to increase endogenous AFF4 mRNA (Fig. 7J) and protein levels (Fig. 7K). Taken together, these results demonstrate that miR-642a-3p directly targets AFF4 and is a strong indirect suppressor of HIV-1, but that its antiviral effect is counteracted by Gag.

## DISCUSSION

Much contention remains over the importance of the components of RNAi in regulating HIV-1 replication, and the existence of virus-encoded RNAi suppressors (36–41). We show here that RNAi mediated by an assortment of exogenous or endogenous siRNAs, shRNAs or miRNAs remains functional in both HIV-1 transfected and infected cells (Fig. 1). We also observed no modification to either localizations of the RISC proteins TRBP, Dicer, Ago2 and DDX6 in HIV-1-producing cells or to their relations with GWBs (Fig. 1D-G), suggesting that the RNAi pathway remains largely unchanged and functional in the presence of HIV-1. Our data corroborate previous findings that while HIV-1 proteins may interfere locally and/or temporally with specific RNAi functions, the virus does not present a net disruptive effect on the RNAi mechanism as a whole (36, 39, 77). These data are compatible with the continued development of antiviral siRNA- or shRNA-based therapies that directly target exposed regions of the HIV-1 genome or indirectly regulate HIV-1 by targeting HDFs. These RNA-based molecules are under development to be used either as drugs or in a gene therapy setting to reach a functional cure against HIV-1 (78–82).

High throughput studies using mass spectrometry are often used to quickly identify large amounts of proteins and the global interactome of specific viral or cellular proteins. One such analysis was performed with all HIV-1 proteins and identified more than 400 protein-protein interactions (29). Proteomic studies using HIV-1 Gag identified 51 main candidate proteins among its interacting proteome (28). Our IF, PLA and IP experiments showed that Gag and Dicer co-localize, are within 40 nm and interact in the absence of RNA in cells (Fig. 1G, 2, 3A-C), supporting the idea of close interaction events between the two proteins. A Gag-Dicer interaction was not observed in the large-scale interactome study (29). However, Dicer was identified as a Gag interacting protein in a second such study, but the interaction was not characterized further (28). Several viral proteins that associate with human Dicer have been found to degrade the endoribonuclease or affect its activity. These effects have been shown for HIV-1 Vpr, Vaccinia virus I7 protease, Dengue virus NS4B, Hepatitis C virus capsid, Zika virus capsid and Ebola virus VP30 and VP35 (31, 83–86). Although this could have been the case for HIV-1 Gag, our results demonstrate that Gag-Dicer interaction neither downregulates Dicer’s expression in cells, nor affects Dicer’s catalytic activity on distinct cellular pre-miRNAs (Fig. 3), suggesting that a functional modification, if any, would be more subtle.

Previous studies have analyzed the differential expression of miRNAs in different HIV-1 infected cells or in people living with HIV-1 (PLWH). Several miRNAs target viral or cellular RNAs to mediate mostly antiviral, but sometimes proviral activities. In the context of HIV- 1 infection, high expression of miRNAs 29a, 28, 125b, 382, 150, 223, 382, 155, 196b and 1290 inhibit viral expression and are associated with maintaining latency in resting primary CD4^+^ T lymphocytes and macrophages (32-35, 73, 87). Specifically, miR-29a is associated with HIV-1 latency and natural resistance by targeting HIV-1 RNA in the overlapping 3’LTR and Nef sequences (48, 71, 88, 89), whereas its expression is decreased during chronic HIV-1 infection (90). Furthermore, miR155 which contributes to HIV-1 latency in resting CD4^+^ T lymphocytes is a biomarker of immune activation in PLWH in extracellular vesicles (32, 73, 91, 92). Some miRNAs target cellular factors, such as miR-198 that targets Cyclin T1 and inhibits viral transcription in monocytes (93), whereas miR-103 and -107 target CCR5 and inhibit viral entry in macrophages (94). While these analyses measure the level of the identified miRNAs, they do not determine their modulation when they are in the RISC. To address this issue, RIP-seq analyses have been performed to identify variations in Ago2-loaded miRNAs in different situations (95–97), but so far, no similar study has identified specific Dicer-loaded miRNAs. These studies combined with our observations of a Dicer-Gag complex formed in cells (Fig. 1-3) suggested that the exploration of miRNAs-bound to Dicer-Gag vs. Dicer alone could have a functional implication in the regulation of HIV-1 replication. Among the selected miRNAs, five were analyzed further either because they were more present on the Dicer-Gag complex or because they were related to HIV-1 infection. Compared to miR-30a-5p and miR-378a-3p, which have an increased expression in HIV-1-transfected cells (Fig. 5) and in some compartments of PLWH (57, 98), RIP-qRT-PCR confirmed that miR-642a-3p, miR-766-5p and miR-766-3p were more associated with Dicer in the presence of Gag (Fig. 5F-H) with no concomitant increase in expression (Fig. 5A-C). This shows for the first time that a viral protein post-transcriptionally regulates specific miRNAs without affecting miRNA biogenesis or RNAi function. Further analyses showed that the collective and shared targets of miR-642a-3p, miR-766-5p and miR-766-3p are functionally enriched in biological processes that affect viral replication either directly or indirectly (Fig. 5K-L). While none of the three noted miRNAs has previously been reported in the context of HIV-1, respiratory syncytial virus produces virus-encoded miRNA seed mimics and miRNA seed sponges that are directly complementary to miR-642a-3p (99), supporting a possible relation between the miRNA and virus regulation in human cells. A downstream analysis on miR-642a-3p, miR-766-5p and miR-766-3p allowed us to confirm their potential gene targets in close relationship with HIV-1 (Table 1, 2 and 3).

Transcriptional elongation is an important regulatory checkpoint in HIV-1 expression and influences whether the virus remains latent or is activated (13, 27, 100). Within the SEC scaffold, AFF4 protein plays a crucial role in triggering and maintaining transcriptional elongation (17, 24–26, 101). In our assays, miR-642a-3p directly regulates AFF4 expression through three main complementary sequences in the mRNA 3’ UTR (Fig. 6). In agreement with AFF4’s known role in HIV-1 transactivation (14, 16, 17, 24–26), this silencing impairs HIV-1 expression (Fig. 7A, B). As miR-642a-3p has not previously been evaluated for its abilities to regulate HIV-1, we compared this miRNA to the previously described anti-HIV-1 miR-29a-3p, which directly targets HIV-1 RNA in the overlapping 3’LTR and Nef coding sequence (48, 72, 102) and to miR-155-5p, which downregulates HIV-1 by directly regulating the viral genome and through the modulation of HDFs (73, 75). Strikingly, miR-642a-3p demonstrated more robust restrictions of HIV-1 expression than did their previously published counterparts (Fig. 7C, D), suggesting a strong involvement of this miRNA in the control of HIV-1 replication.

Because miR-642a-3p is more present on Dicer-Gag than on Dicer alone (Fig. 4D, 5F), the next question was to assess the role of Gag in miR-642a-3p’s regulation of AFF4 expression. Our results show that miR-642a-3p’s regulation of AFF4 and downstream HIV-1 expression is impaired in the presence of Gag but not in its absence (Fig. 7A, F, G), indicating that Gag counteracts the effects of this miRNA. In agreement with these experiments, Gag increases the concentration of endogenous AFF4 in cells that are not overexpressing the miRNA (Fig. 7H-J). These data strongly suggest that upon its interaction with Dicer, Gag prevents miR-642a-3p’s function allowing an increase of AFF4 expression. Some similarity can be made with cellular proteins that share specific features with Gag like Lin28a. Indeed, Lin28a is a cellular RNA-binding protein that interferes selectively with the biogenesis of miR-let7 by binding to the miRNA’s terminal loop during Dicer processing (103, 104). Lin28a shares homology with a conserved Zinc-knuckle domain in Gag’s nucleocapsid product, which recognizes nucleotide sequences GGAG and GGUG (105). Interestingly, a GGAG motif is present in mature miR-642a-3p and may be a possible recognition motif that facilitates or fastens the interaction with Dicer-Gag. Further investigations into the Dicer-Gag interaction using proteomic, transcriptomic and structural biological techniques will be needed to fully elucidate the interaction partners and molecular events involved in the post-translational modulation of miR-642a-3p by Gag.

In summary, our results support the paradigm that HIV-1 does not induce major dysfunctions on miRNA processing or RNAi mechanism. Nevertheless, Gag can interact with Dicer and induce subtle post-transcriptional differences in the occupancy of specific antiviral miRNAs on Dicer. Among three miRNAs retained on Dicer, miR-642a-3p was found to regulate AFF4, and by extension HIV-1. This regulation was impeded by Gag, suggesting that the viral structural protein sequesters miR-642a-3p-loaded Dicer complexes, thereby hindering or preventing Ago loading (Fig. 8). Collectively, these results support the existence of a novel post-transcriptional mechanism by which a mammalian virus selectively modulates the cellular regulatory environment, which results in the enhancement and auto-amplification of viral expression. Counteracting this mechanism could increase HIV-1 silencing and contribute to its long-term latency in a blocking strategy. On a larger view, similar mechanisms may occur with other pathogens, during normal cell function, development and various diseases (106, 107).

**Figure 8:**
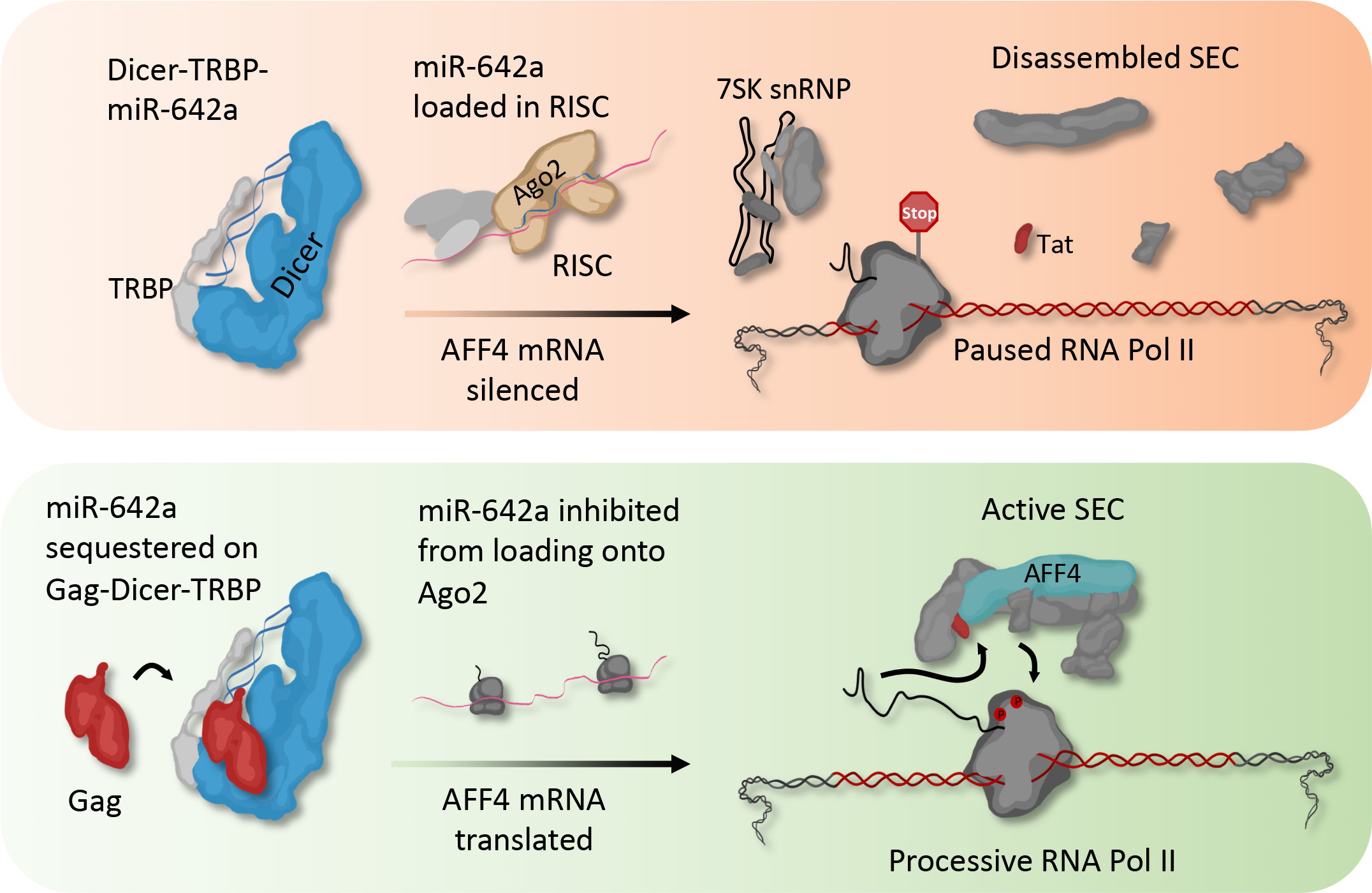
Schematic representation of miRNA retention on the Gag-Dicer complex inducing HIV-1 transcription elongation. (Top) In the absence of HIV-1, miR-642a-3p is loaded on Dicer-TRBP, which recruits Ago2 to form the RNA-induced silencing complex (RISC). miR-642a-3p targets and silences AFF4 mRNA, resulting in a poor assembly of the super elongation complex (SEC) and paused RNA polymerase (Pol) II. (Bottom) In the presence of HIV-1, Gag binds to Dicer, which induces the sequestration of miR-642a- 3p and prevents its loading on the RISC. AFF4 mRNA is translated, which results in the formation of an active SEC including AFF4 and a processive RNA Pol II.

## MATERIALS AND METHODS

### Cell lines and growth conditions

HeLa cells were originated from a human female cervix whereas HEK 293T cells were from a human female embryonic kidney. HeLa (CCL-2) and HEK 293T (CRL-11268) adherent cells were originally obtained by the American Type Culture Collection (ATCC). Both cell lines were grown and maintained in Dulbecco’s modified Eagle’s medium (DMEM) (Hyclone) supplemented with 10% fetal bovine serum (FBS) (Hyclone), 50 μg/mL streptomycin, and 50 U/mL penicillin (Gibco) at 37°C and 5% CO2 in a humidified incubator.

### Plasmid constructions

The HIV-1 molecular clone pNL4-3 was obtained from NIH AIDS Reference and Reagent Program (ARRP), originally described (120). pNL-XX (pNL4-3 with no Gag expression) was kindly provided by Dr. David Ott (NCI Frederick, MD). pNL-XX contains a six nucleotide mutation in *gag*’s initiation codon that produces an amber nonsense codon (TAG) as well as an additional nonsense mutation within CA to ensure Gag is not expressed, as previously described (76). GAPDH-ProLabel, pSIREN-shRNA-LucIRR and pSIREN-shRNA GAPDHHB were purchased from Clontech. pCDNA3.1 was purchased from Invitrogen. pcDNA3h/TO-HA-AFF4 was obtained from Dr. Qiang Zhou (University of California, Berkeley) (70). pCI-Neo-Flag (pFlag) and pCI-Neo-Flag-Gag (Flag-Gag) were previously described (121). The shRNA against EGFP and the EGFP-clet7 plasmids were previously described (36, 42). GST-Dicer was donated by Dr. W. Filipowicz (5). GST-Dicer was used as a Dicer gene template to construct pCMV2-HA-Dicer (HA-Dicer). The gene was amplified by PCR using 5’AGCGGTCGACCATGAAAAGCCCTGCTTTG3’ as a forward primer and 5’ATATAGCGGCCGCTCAGCTATTGGGAACCTG3’ as a reverse primer. These PCR products were digested by SalI and NotI and subcloned in frame into pCMV2-HA (Clontech) cut by the same enzymes. We previously described the generation of psiRNA-U6GFP::Zeo and the nonsense negative control psiRNA-U6GFP::Zeo-shNS from the U6 RNA Polymerase III promoter in pSIREN-shuttle (Clontech) and psiRNA- 7SKGFP::Zeo (InvivoGen) (122, 123). To generate psiRNA-U6GFP::Zeo-shAFF4, the following complementary oligonucleotides were annealed (1.25 µM each in 75 mM NaCl, 40 µL, 2 min at 80 °C, cooled over 1.5 h to 37 °C) as in (124). Sense shAFF4:

5’ACCTCGCACCAGTCTAAATCTATGTTCTCGAGAACATAGATTTAGACTGGTGCTTT

3’ Antisense shAFF4: 5’CAAAAAAGCACCAGTCTAAATCTATGTTCTCGAGAACATAGATTTAGACTGGTGCG

3’

The annealed double-stranded inserts were then ligated into Bbs1-digested psiRNA- U6GFP::Zeo resulting in the plasmid psiRNA-U6GFP::Zeo-shAFF4.

The pEGFP-C1 plasmids including AFF4 3’UTR sites “1-2-3” were generated using pEGFP-C1 (Clontech). The following complementary oligonucleotides were annealed in respective pairs as above (1.25 µM each in 75 mM NaCl, 40 µL, 2 min at 80 °C, cooled slowly to 37 °C).

Sense pEGFP-C1-1-2-3: 5’TCGAGCATAATGATATGTACTATGAAATGTGTCTGATTATATTTTCTCTTTAAAACTGTGTCAATTTCCCCCCTCCCTCCTCAATAGGTGTCCGGTAC3’

Antisense pEGFP-C1-1-2-3: 5’CGGACACCTATTGAGGAGGGAGGGGGGAAATTGACACAGTTTTAAAGAGAAAATATAATCAGACACATTTCATAGTACATATCATTATGC3’

The annealed double-stranded inserts were then ligated into XhoI- and KpnI-digested pEGFP-C1. The resulting constructs were confirmed by sequencing using a primer located in the EGFP ORF: 5’TCACATGGTCCTGCTGGAGTT3’.

### Cell transfection

For Western blots, RNA extraction and immunoprecipitations (IP), 10 cm culture plates were seeded with 2.6 x 10^6^ cells or 6-well plates were seeded with 5 x 10^5^ cells/well 12- 15 h prior to transfection, whereas 5 x 10^4^ cells/well were seeded in 96-well plates 24 h prior to transfection for indirect immunofluorescence (IF) assays. If not otherwise specified in figure legends, cells were fixed or lysed 48 h after transfection. Depending on the plasmid, cells were transfected with 0.25-1 μg of DNA per millilitre of cell media. DNA was introduced into cells using polyethyleneimine MAX 40K (PEI MAX) (Polysciences) at a ratio of 3:1 PEI:total DNA or TransIT-LT1 (Mirus) at a ratio of 3:1 TransIT:DNA. mirVana® miRNA mimics and anti-miR™ miRNA inhibitors (Thermo Fisher) were transfected according to the manufacturer’s protocols using Lipofectamine RNAi MAX (Thermo Fisher) 48 h prior to transfection by other plasmids using PEI MAX.

### Antisera against MCEF/AFF4 (anti-AFF4-7D)

The cDNA for AFF4 (also known as MCEF) amino acids 1-715 was cloned into pet-6HIS- 11d plasmid (Novagen) and expressed in E. coli BL21 cells. The protein was purified and injected into rabbits as previously described (66). A pre-bleed was taken for controls followed by bleeds at different times post-injection. After a primary injection on day 1 (200 μg), boosters were given on days 21, 45, 66, 86, 107 and 128 (200 μg each). Several sera were tested and for the experiments in this paper, we used the polyclonal antisera collected at day 146 (7D). Serum 7D was validated using overexpressed HA-AFF4 (70) compared to endogenous AFF4 protein (Fig. S4).

### Immunoblotting

Cells were washed twice with phosphate-buffered saline (PBS) (Wisent) and lysed in cold immunoblotting lysis buffer [50 mM Tris-HCl pH 7.4, 150 mM NaCl, 5 mM EDTA pH 8.0, 10% Glycerol, 1% IGEPAL® CA-630 (Sigma)] with phosphatase and protease inhibitor cocktails (Roche) or cold radioimmunoprecipitation assay (RIPA) buffer (50 mM Tris, pH 8.0, 150 mM NaCl, 1% IGEPAL^®^ CA-630, 5 mM EDTA pH 8.0, 0.1% sodium deoxycholate, 0.5% SDS) with phosphatase and protease inhibitor cocktails (Roche). The lysates were then freeze-thawed three times between liquid nitrogen and ice. RIPA lysates were incubated with 1 μL Benzonase and 1 μL MgCl2 for 30 min at room temperature. All samples were then centrifuged for 30 min at 13000 rpm on a tabletop centrifuge to remove debris. Proteins in preserved supernatants were quantified by Bradford assay (Bio-Rad). 35-125 μg of protein mixed with Laemmli sample buffer were incubated for 5 min at 95 °C. Proteins were separated by sodium dodecyl sulfate polyacrylamide gel electrophoresis (SDS-PAGE) and wet-transferred overnight at 150 mA to Hybond nitrocellulose membranes (Bio-Rad) as described (125) or fast-transferred using TransBlot Turbo (Bio-Rad) using preprogrammed Turbo protocols for membranes probing lower molecular weight proteins (126, 127). Membranes were blocked for 1 h with 5% milk in 0.1% Tris-buffered saline with tween 20 (TBS-T) followed by three 5-minute washes with TBS-T. Membranes were then incubated overnight at 4 °C with anti-GST antibody (GE Healthcare) at a 1/1000 dilution, anti-HIV-1 p24 (183-H12-5C) antibody (National Institute of Health AIDS reagent program) at a 1/10;000 dilution, anti-AFF4 (7D) antibody at a 1/500 dilution, anti-HA (Sigma H6908) anti-GAPDH antibody (Clontech) at a 1/2500 dilution, anti-actin antibody (Chemicon) at a 1/10000 dilution, or anti-Dicer 13D6 antibody (Abcam) at a 1/1000 dilution. One-hour blots were performed using anti-EGFP antibody (Santa Cruz Biotechnology) at a 1/1000 dilution, anti-HIV-1 p24 (183-H12-5C) at a 1/1000 dilution, anti-HIV-1 Reverse Transcriptase (RT) antibody (from Dr. Lawrence Kleiman at Lady Davis Institute) at a 1/1000 dilution, anti-GAPDH antibody (Santa Cruz Biotechnology) at a 1/1000 dilution. After three 5-minute washes with TBS-T, membranes were incubated with horseradish peroxidase-conjugated secondary antibodies (Amersham, SeraCare, Millipore or Rockland Immunochemicals) at a 1/5000 dilution for goat anti-rabbit, 1/5000 dilution for mouse anti-rabbit light chain, and 1/3000 for goat anti-mouse. After three 5-minute further washes with TBS-T, the bands were visualized with Western Lightning Plus-ECL reagent (Perkin-Elmer) or Pierce^TM^ ECL Western Blotting substrates (Thermo Fisher). Western blots are representative of at least three independent replicates.

### Immunofluorescence and imaging

HeLa cells transfected with pNL4-3 were subjected to IF at different time points (24, 48, 36 and 72 h). Cells were washed with PBS (Wisent) and fixed with 4% formaldehyde / 4% sucrose for 20 min. Cells were then washed once with PBS and incubated with 0.1 M glycine for 20 min. Cells were washed once again with PBS and permeabilized with 0.2% Triton X-100 for 5 min. After permeabilization, cells were washed with PBS and blocked with bovine serum albumin (BSA) 3%. Cells were incubated with the primary antibodies for 1 h at 37 ℃ using mouse monoclonal anti-p24 (183-H12-5C) (128) at a dilution of 1/400 or 1/1000, rabbit anti-p24 (from Dr. L. Kleiman) at a 1/1000 dilution, rabbit anti- Rck/p54/DDX6 (Bethyl labs) at a 1/1000 dilution, mouse anti-Ago2 (Wako 4G8) at a 1/100 dilution, rabbit anti-TRBP673 (129) at a 1/250 dilution, rabbit anti-Dicer 349 (5) at a 1/250 dilution, mouse monoclonal anti-Dicer 13D6 antibody (Abcam) at a 1/500 dilution and human autoimmune serum 18033 (130) at a 1/5000 dilution. Cells were washed 5 times for 5 min with a rinsing solution (PBS, tween 20 0.3%, BSA 0.1 %), and were then incubated with Alexa 488, 546 or 647 (Invitrogen) secondary antibodies in total darkness for 1 h on a rocking platform. Cells were rinsed again in the dark 5 times for 5 min. Coverslips were then mounted with Immu-Mount (Thermo-electron corporation). Imaging was performed on a Zeiss Pascal confocal laser scanning microscope (LSM) using a 65X objective. Image acquisition was carried out using LSM AIM Pascal acquisition software. Image processing and analysis were performed using Imaris software (version 8.4.1 Bitplane Andor) and Pearson’s R-values (no threshold) were calculated using Fiji (131).

Imaging experiments were performed at least three times.

### Co-immunoprecipitations (Co-IPs)

HEK 293T cells were transfected with GST-Dicer, HA-Dicer and/or Flag-Gag. After 48 h, cells were washed once with PBS and lysed with cold immunoblotting lysis buffer as above. Total proteins were quantified by Bradford assay (Bio-Rad). For each condition, 1000 µg of protein extracts were cleared with agarose protein A beads (Millipore) at 4 ℃ for 1.5 h on a rocking platform. Samples were centrifuged at 8000 rpm for 1 min in a tabletop centrifuge and the supernatant was collected. 2 µl of either anti-GST (GE Healthcare), anti-HA (Sigma H6908), anti-Dicer 13D6 (Abcam) or isotype (Invitrogen) were added to the supernatant and incubated with constant shaking at 4 ℃ overnight. 30 µl of Dynabeads protein A (Thermo Fisher) or Sepharose protein G (Sigma) were added and further incubated with shaking for 3 h at room temperature. Beads were pelleted by centrifugation or magnetic separation and washed three times for 5 min with lysis buffer. All supernatants were discarded and 30 µl of Laemmli sample buffer was added and incubated for 5 min at 95 °C. Samples were loaded for SDS-PAGE analysis and analyzed by immunoblotting.

### *In situ* protein-protein interaction assay

HeLa cells were transfected with Flag-Gag or pNL4-3 for 48 h, followed by *in situ* proximity ligation assay using the DUOLINK II In Situ kit (Duolink) according to the manufacturer’s protocol. PLA is based on two primary antibodies from different species that target different proteins of interest, followed by secondary antibodies with oligonucleotide tails attached to their Fc regions. If the primary antibodies’ targets are within 30 nm of each other, these oligonucleotides permit the rolling circle amplification of a set of complementary detection oligonucleotides. Labeled oligonucleotides are then added to probe for the detection oligo amplicons, emitting a fluorescent signal in the event of a <30 nm interaction which is visualized as a discrete fluorescent dot (132). PLA conditions were previously described (133, 134). Mouse monoclonal anti-Dicer 13D6 antibody was used at a 1/500 dilution and rabbit polyclonal anti-HIV-1 SF2 p24 (National Institute of Health AIDS reagent program) was used at a 1/400 dilution. The antibodies were detected using DuoLink II Detection Reagent Red, Duolink II PLA Probe Anti-Mouse MINUS, and DuoLink II PLA Probe anti-rabbit PLUS. Imaging was performed on a confocal LSM Leica DM1600B equipped with a WaveFX spinning disk confocal head (Quorum Technologies). Data analysis was performed as described using Imaris software (version 8.4.1 Bitplane Andor) (134, 135). Statistical analyses for Dicer-Gag interaction were performed using Graph-Pad Prism V6 witih a unpaired Mann-Whitney u tests. For quantification of nuclear colocalization intensity, raw .liff files were exported by the Volocity software (Perkin Elmer) for import into Imaris and ImarisColoc software v.8.4.1 (Bitplane/Andor) used for generation of new colocalization channels of nuclear DAPI-PLA, and .csv exports of quantitative measurements of mean signal intensity values used for downstream data harmonizing and statistical analyses using Excel (Microsoft) and GraphPad Prism v6 (unpaired *t*-tests).

### Dicer catalytic activity in the presence of HIV-1 Gag

The evaluation of Dicer catalytic activity was performed according to (47). Briefly, we transcribed T7 promoter-pre-Let7c 5’GCTCCUUGGUUUGCTUGUUGGTTGTUCUGTTUUCTCCCUGGGTGTUUCTCTUUUCCUTUCUUCCTUCTUCCTCUUCCCGGUTGCCCTATAGTGTGAGTCGTATTA 3’ and T7-promoter-pre-miR29a 5’ATAACCGATTTCAGATGGTGCTAGAAAATTATATTGACTCTGAACACCAAAAGAAATCAGTCCCTATAGTGAGTCGTATTA3’ using the T7 high yield RNA synthesis kit (BioLabs) with α ^32^P UTP (Perkin Elmer) at 800-6000 Ci/mmol ≥ 10 mCi/mL. Pre-miRNAs were cleared from the DNA template using DNase I (QIAGEN) for 15 min and 20 µl of GLB II (Thermo Fisher Scientific), mixed and loaded on a denaturing 10% polyacrylamide gel (19:1)/ 7 M urea. Pre-miRNAs were purified from the gel followed by heating and slow cooling for proper folding. 45,000 cpm of labeled RNA were incubated with Dicer IP complexes attached to Sepharose protein G beads for 1 h at 37 ℃. RNA was extracted and concentrated by TRizol and ethanol precipitated with Glycogen and yeast tRNA (Thermo Fisher Scientific). Products were subjected to denaturing PAGE and visualized in X-ray films after incubation at -80 ℃ for 48 h.

### RNA immunoprecipitation sequencing (RIP-seq) for small RNA detection

RIP-seq experiments were based on (136) with modifications. Briefly, HEK 293T cells were transfected with HA-Dicer/pFlag, HA-Dicer/Flag-Gag, HA/Flag-Gag (negative control). After 48 h, cells were washed once with PBS and lysed with RNA extraction buffer (150 mM NaCl, 50 mM Tris, pH 7.5, 5 mM EDTA, 0.5% Nonidet P-40, 0.5% of RNaseOUT (Invitrogen), 0.2% of Vanadyl (NEB), 100mM of DTT (Invitrogen) protease and phosphatase inhibitor cocktail (Roche)). 30 µl of agarose protein A beads (Millipore) washed with RNA extraction buffer were added to 1 mg of protein lysate for a 1.5 h incubation at 4 ℃ on a rocking platform. After centrifugation, the supernatant was incubated with 50 µl of 1% BSA blocked Pierce anti-HA magnetic beads (Thermo Fisher) or 1% BSA blocked Pierce NHS-activated magnetic beads (Thermo Fisher) conjugated with a mouse monoclonal IgG1 isotype antibody (Invitrogen) overnight at 4 ℃ in constant rotation. The following day, samples were magnetically separated and the flow-through was discarded. The magnetic beads were washed three times with RNA extraction buffer and separated in two samples for protein and RNA analysis. For protein analysis, 30 µl of Laemmli sample buffer was added to the beads and incubated for 5 min at 95 °C. RNA extraction was performed on the second sample of the beads using Trizol reagent and the miRNeasy mini kit (QIAGEN). An RNase-free DNase (QIAGEN) was used according to kit protocols in the miRNeasy columns to eliminate possible DNA contamination. Libraries were synthesized from 300 ng of total RNA with the New England BioLabs NEBNext^®^ Small RNA Library Prep Kit for Illumina^®^. Libraries were run in an Illumina HiSeq2500 SR50 sequencing lane at Génome Québec, Canada.

### Small non-coding RNA analysis

RNA bioinformatics were performed at the Canadian Centre for Computational Genomics (McGill University). Trimming and clipping were performed using Trimmomatic (137).

Illumina sequencing adapters were removed and reads were trimmed from their 3’ end to have a minimum phred score of 30. The filtered reads were aligned to *Homo sapiens* assembly GRCh37 using STAR (138). All readset BAM files were merged into a single global BAM file using Picard (139). All control samples (HA/Flag-Gag and the isotype antibody sample) were merged into a single file as well to use with peak-calling software. Peaks were called using Model-based Analysis of ChIP-Seq (MACS) v2 software (49). The mfold parameter used in the MACS model building step was estimated from a peak enrichment diagnosis run. The estimated mfold lower bound was 5 and the estimated upper bound was 50. Nmodel and extsize were adjusted to better reflect the conditions of the experiment. The peak calling strategy was based on pooled samples (differences between the samples could be compensated during peak calling, producing a smaller but more robust set of peaks) and individual samples to increase the sensitivity.

All called peakset files were annotated using annotatePeaks.pl from Homer (140). The information used to annotate the peaks came from Ensembl 75 (141), which includes information on gene transcripts as well as the positions of 9,459 miRNA, 5,783 snRNAs, and 54,912 ncRNAs. To determine differences between peaks from both experimental conditions, a differential binding analysis was carried out using the R package DiffBind (50). The purpose of this package is to find overlapping peaks between the samples, merging the peak sets and then counting reads in the overlapping intervals of the peak sets. The differential binding analysis is carried on by an exploratory step to determine the potential occupancy of a specific locus (occupancy analysis). To corroborate the annotation of miRNAs produced by Diffbind-occupancy analysis, miRNA sequences were aligned to *Homo sapiens* GRCh37 (hg19) with IGV software (52–54). miRNAs with high coverage in both strands (5p and 3p) during the alignment were considered as two independent miRNAs. The latter generated a total of 22 miRNAs that were plotted by sample in a heatmap (log2) made in R pheatmap. Next, sequences of the 22 miRNAs were filtered out on miRbase using their integrated web-based BLASTN search tool (55, 56, 142). Only those miRNAs that matched exactly with sequences and IDs displayed on miRBase were further analyzed and normalized to the total reads. Quantitative differences of filtered miRNAs were further normalized to the coverage of HA/Flag-Gag and isotype controls. A two-way ANOVA with Sidak’s multiple comparisons test was performed on the final normalized set of miRNAs from each group.

### RIP and Reverse Transcription Quantitative Polymerase Chain Reaction (RT-qPCR)

For RIP RT-qPCR experiments, 2.6 x10^6^ HEK 293T cells were co-transfected with a total of 9 µg of HA-Dicer and Flag-Gag. Cells were collected 48 h post-transfection, washed once with PBS and lysed with RNA extraction buffer for 15 min at 4 ℃. Lysates were centrifuged on a tabletop centrifuge at 13,200 rpm at 4 ℃ for 15 min. 2.2 mg of protein supernatants were used for the IP and 0.022 mg (1%) was used as the input. 30 µl of agarose protein A beads (Millipore) were washed once with RNA extraction buffer and diluted with 25 µl of RNA extraction buffer before being combined with the protein supernatants for 1.5 h at 4 ℃ with constant agitation. Samples were centrifuged at 8000 rpm for 1 min, and supernatants were incubated with 2 µl of the anti-HA antibody or 1 µl of control rabbit serum overnight. 50 µl of pre-blocked Dynabeads protein A with 1% BSA RNA extraction buffer were added to the beads and incubated with constant agitation for 3 h at 4 ℃. Supernatants were discarded and the beads were washed three times with RNA extraction buffer (without RNaseOUT) for 5 min with a magnetic stand. After the final wash, 700 µl of TRIzol reagent and 140 µl of chloroform were added to the beads. To extract the RNA, QIAGEN’s miRNeasy mini kit was used according to the manufacturer’s protocol. QIAGEN’s RNase-free DNase was loaded onto the RNeasy columns to avoid any DNA contamination. Due to the low abundance of RNAs in IPs, we used QIAGEN’s isopropanol protocol for a better recovery of RNAs. The remaining RNA was diluted from columns in 60 µl of ultrapure distilled water. Immunoprecipitated RNAs were concentrated by precipitating the eluted samples with 50 µl of ammonium acetate, 5 µl of Glycogen and 700 µl of ice-cold ethanol 100% at -80 ℃ for one hour, followed by 30 min of centrifugation at 13,200 rpm on a tabletop centrifuge. The pellet was washed once with ice-cold ethanol 70%, dried and diluted in 11.5 µl of ultrapure distilled water. 1.5 µl were used to measure the RNA in a Spectrophotometer/Fluorometer (DeNovix, DS-11 FX+). RNA concentrations from IPs were 40-60 ng/µl for HA-Dicer/pFlag and 50-75 ng/µl for HA- Dicer/Flag-Gag. cDNA synthesis, miRNA detection and amplification were performed as in (59) with at least three independent experiments and at least three technical replicates for each independent experiment. Table 4 shows the primers to track specific miRNAs.

**Table 4.**
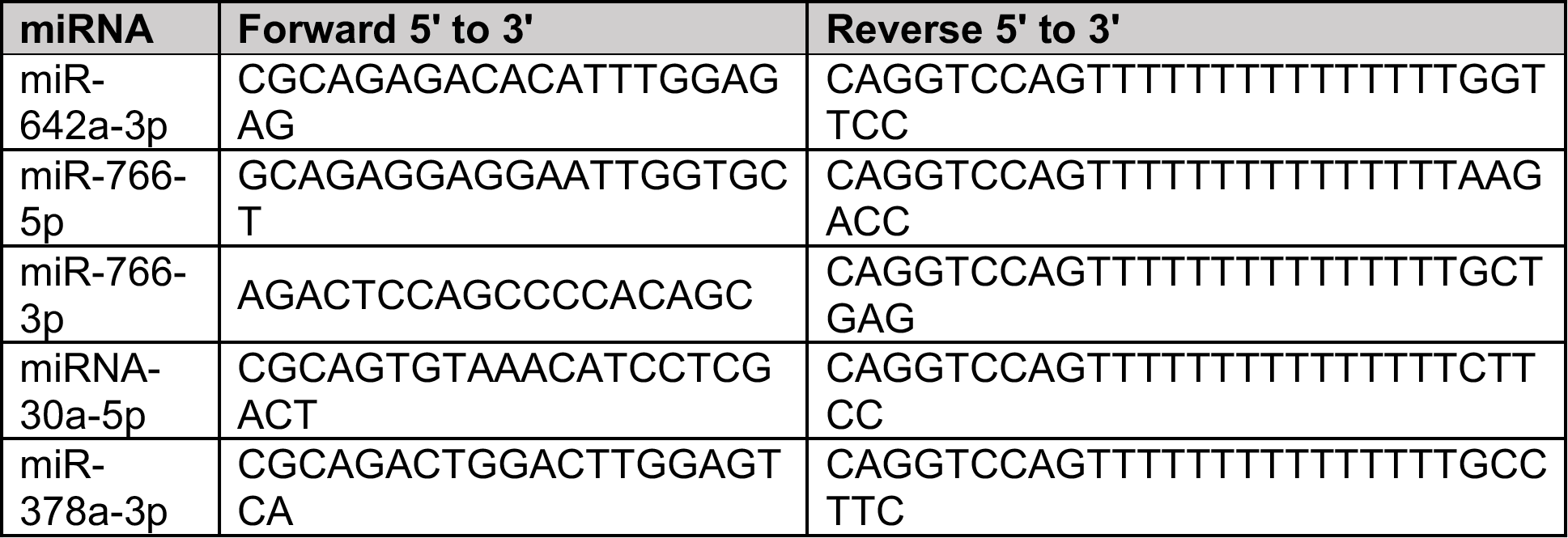
Primer sequences to evaluate selected miRNAs.

Quantitative PCR data acquisition and analysis were performed using Bio-Rad CFX96 and CFX Maestro software, respectively (143, 144). Actin was used as a housekeeping control to calculate ΔΔCq and percent input [100*2^ (Adjusted input - Ct (IP)] for the abundance of miRNAs in IPs. The calculated abundance was transformed from percentage to fold change and *t*-tests (n=3) were carried out in Graph-Pad Prism V6.

### miRNA Network Generation, Target Prediction and Enrichment

miRNAs identified in RIP-seq and RT-qPCR experiments to be associated with Dicer in Flag-Gag transfected cells were used as queries for target entries in miRTarBase v.8.0 and TarBase v.8.0. All entries in these databases (low- and high-scoring) for targets associated with these human miRNAs were used for visualization and enrichment analysis. miRNA-target interactions were visualized using MirNet 2.0 (145). Enrichments of these targets were quantified using a Fisher’s exact test for Biological Processes (BP) GO terms, in which experimentally validated targets were queried against the background of human miRNA targets listed in miRTarBase v.8.0 and TarBase v.8.0 databases. The Fisher’s exact test with Bonferroni correction for multiple testing was performed on PANTHER (146) according to the 2021-02-01 release of the Gene Ontology (GO) Resource. miRNA targets were predicted using online prediction databases miRDB and TargetScan release 8.0, which use distinct data-informed algorithms to predict and score miRNA-target pairings (60, 68).

### RT-qPCRs

Cells incubated in 6-well plates for 48 h following transfection were lysed using 750 μl of TRIzol (Life Technologies). RNA extraction was performed using the miRNeasy® mini kit (QIAGEN) according to the manufacturer’s protocol, including digestion of contaminating DNA using an RNase-free DNase set (QIAGEN). RNA was diluted from the columns in ultrapure distilled water. 1 μg of total RNA was used to synthesize cDNA using Oligo(dT)12-18 and SuperScript™ II Reverse Transcriptase (Thermo Fisher) according to the manufacturer’s protocol. qPCR data acquisition and analysis were performed using Bio-Rad CFX96, primers in Table 5 and CFX Maestro software, respectively, using ABM’s 2X BrightGreen qPCR MasterMix according to the manufacturer’s protocol with at least three technical replicates for each biological replicate. Gene specific mRNA expression is normalized to actin. Actin primers are described in this manuscript and elsewhere (147).

**Table 5.**
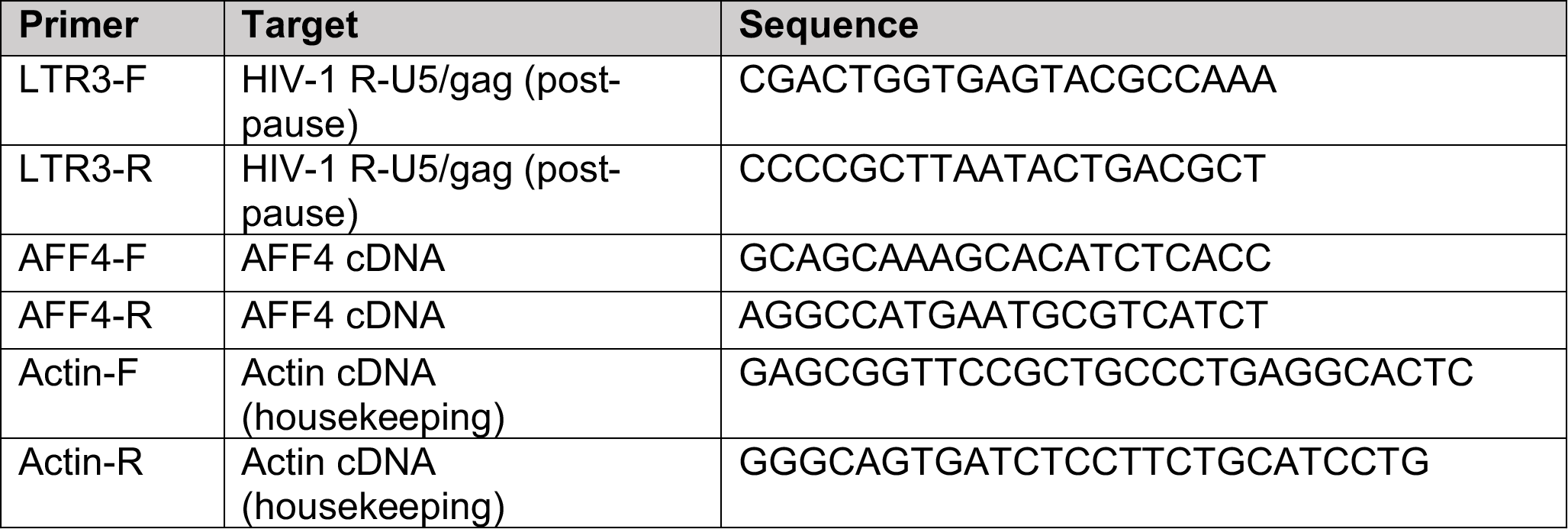
List of primers used in qPCR analyses

Statistical tests were carried out in Graph-Pad Prism V6. Error bars in graphs represent the ±SEM of three independent experiments and *t-*tests were performed to assess significance.

### Water-soluble tetrazolium salt 1 (WST-1) assay

The WST-1 assay was used to check cell viability in response to treatment by different antiviral miRNA mimics (122, 143). This colorimetric assay measures the activity of NAD(P)H-dependent cellular oxidoreductase enzymes in reducing the WST-1 to the insoluble dye Formazan, which is taken as a quantifiable correlate for proliferating live cell count. HEK 293T cells were incubated in 96-well plates with miRNAs for 72 h, after which they were exposed to WST-1 according to the manufacturer’s protocol (Roche). The Formazan absorbance of each sample was measured at a 450 nm wavelength using a Synergy 4 (BioTek) microplate spectrophotometer, while a control reading at 690 nm was used to control for microplate imperfections. Absorbance values were normalized to the nonsense miRNA control.

### Quantification and statistical analysis

All experiments were performed in triplicates unless otherwise indicated. Unless otherwise stated, non-grouped analyses were performed using Students’ *t*-tests and grouped data were analyzed by one-way ANOVA with correction for multiple comparisons, both performed in GraphPad Prism v8 or in Microsoft Excel. Error bars denote ±SEM or SD as indicated in figure legends. ns, p > 0.05. ∗, p < 0.05. ∗∗, p < 0.01. ∗∗∗, p < 0.001, ∗∗∗∗, p < 0.0001 Statistical parameters for each experiment are provided in figure legends. Fig 1D, bar graph, unpaired Mann-Whitney test p24+ve versus p24-ve, p=0.7 (ns) Fig 2C, boxplot in log2, unpaired Mann-Whitney test: pcDNA3.1 versus pNL4-3, p < 0.0001. pFlag versus Flag-Gag p < 0.0001 Fig 2D, bar graph, unpaired *t*-test: pcDNA3.1 versus pNL4-3, p < 0.05. pFlag versus Flag- Gag, p < 0.05 Fig 4D, bar graph, two way ANOVA, Sidak’s multiple comparison test: HA-Dicer/pFlag versus HA-Dicer/Flag-Gag, p>0.05 except for miR642a-3p, p<0.05 Fig 5A, bar graph, unpaired *t*-test: miR-642a-3p concentration (HA-Dicer/pFlag versus HA-Dicer/Flag-Gag) p>0.05 Fig 5B, bar graph, unpaired *t*-test: miR-766-5p concentration (HA-Dicer/pFlag versus HA- Dicer/Flag-Gag) p>0.05 Fig 5C, bar graph, unpaired *t*-test: miR-766-3p concentration (HA-Dicer/pFlag versus HA- Dicer/Flag-Gag) p>0.05 Fig 5D, bar graph, unpaired *t*-test: miR-30a-5p concentration (HA-Dicer/pFlag versus HA- Dicer/Flag-Gag) p>0.05 Fig 5E, bar graph, unpaired *t*-test: miR-378a-3p concentration (HA-Dicer/pFlag versus HA-Dicer/Flag-Gag) p>0.05 Fig 5F, bar graph, unpaired *t*-test: miR-642a-3p fold increase (HA-Dicer/pFlag versus HA- Dicer/Flag-Gag) p< 0.001 Fig 5G, bar graph, unpaired *t*-test: miR-766-5p fold increase (HA-Dicer/pFlag versus HA- Dicer/Flag-Gag) p< 0.05 Fig 5H, bar graph, unpaired *t*-test: miR-766-3p fold increase (HA-Dicer/pFlag versus HA- Dicer/Flag-Gag) p< 0.05 Fig 5I, bar graph, unpaired *t*-test: miR-30a-5p fold increase (HA-Dicer/pFlag versus HA- Dicer/Flag-Gag) p< 0.05 Fig 5J, bar graph, unpaired *t*-test: miR-378a-3p fold increase (HA-Dicer/pFlag versus HA- Dicer/Flag-Gag) p< 0.01 Fig 6C, bar graph, unpaired *t*-test: shNS versus shAFF4 p < 0.0001. miR-NS versus miR- 642a-3p p < 0.001. Fig 7A, bar graph, unpaired *t*-test: AFF4transcription (miR-NS versus miR-642a-3p) p<0.05. HIV-1 expression (miR-NS versus miR-642a-3p) p<0.05 Fig 7C, bar graph, two way ANOVA, Tukey’s multiple comparisons test for HIV-1 Gag (p55, p41, p24): shNS vs shAFF4 (p=0.0008), miR-NS versus miR-642a-3p (p= 0.0016), antimiR-NS versus antimiR-642a-3p (p= 0.9685). Fig 7E, bar graph, two way ANOVA, Dunnett’s multiple comparisons test for HIV-1 Gag (p55, p41, p24): miR-NS versus miR-29a-3p (p < 0.0001), miR-155-5p (p < 0.0001), miR- 642a-3p (p < 0.0001). Fig 7G, bar graph, unpaired *t*-test: AFF4 transcription-HIV-1 no Gag (miR-NS versus miR- 642a-3p) p<0.01. HIV-1 expression-HIV-1 no Gag (miR-NS versus miR-642a-3p) p<0.05. Fig 7J, bar graph, a one way ANOVA (p=0.0046) and a subsequent Dunnett’s multiple comparisons test p(untransfected/Flag-Gag)=0.0262, p(pFlag/Flag-Gag)=0.0685 (ns).

### Data and code availability

Sequencing data associated with RIP-seq analysis have been deposited in GEO (GEO: GSE201502 - RNA immunoprecipitation sequencing (RIP-seq) of human Dicer- bound miRNAs in HIV-1 Gag replicating cells).

## SUPPLEMENTAL MATERIAL (attached)

Supplemental material is available online only.

## ACKNOWLEDGMENTS

We thank Drs. Eric Lécuyer (IRCM, Montréal, QC, Canada) and Pascale Legault (University of Montréal) for valuable discussions, James Saliba (LDI and McGill University) for helping in the qRT-PCR analysis, Anne Monette (LDI) for her direction in confocal microscopy analysis and Valérie Le Sage (LDI) for help in PLA assays. We also thank José Héctor Gálvez López and François Lefebvre (Canadian Centre for Computational Genomics, Montréal) for the pre-processing and bioinformatic analysis of RIP-seq reads. We are grateful to Drs. Witold Filipowicz for providing GST-Dicer and anti- Dicer 349 antibody, Lawrence Kleiman for anti-p24 and anti-RT antibodies, Marvin Fritzler for the human serum 18033, David Ott for pNL-XX and Qiang Zhou for HA-AFF4. We are also thankful for expert technical assistance from Meijuan Niu (LDI). The following reagents were obtained through the NIH AIDS Reagent Program, Division of AIDS, NIAID, NIH: anti-HIV-1 p24 monoclonal antibody (183-H12-5C) and HIV-1 strain NL4-3 infectious molecular clone (pNL4-3).

This study was supported by grants from the Canadian Institutes of Health Research (CIHR) (MOP-136897 to AG), (PJT-148704 to AG and RJS) and (MOP-56974 to AJM), by the Canadian HIV Cure Enterprise Team Grant HIG-133050 (to AG and AJM) from the CIHR in partnership with the Canadian Foundation for AIDS Research (CanFAR) and the International AIDS Society (IAS), and by a grant from the France-Canada Research fund (to AG and BM). SPAL was supported by a Doctoral fellowship from the Consejo Nacional de Ciencia y Tecnologia (CONACYT) (Mexico). ORSD was supported by graduate scholarships from the CIHR, the Fonds de Recherche du Québec - Santé (FRQS) and from the McGill University Division of Experimental Medicine. RJS was a recipient of a post-doctoral fellowship from the Richard and Edith Strauss Canadian Foundation through the McGill University Department of Medicine.

## AUTHOR CONTRIBUTIONS

Conceptualization: SPAL, AM and AG; methodology and investigation: SPAL, ORSD, RJS, SMD, AD, MT and MCE; formal analysis: SPAL, ORSD, RJS and MT; writing original draft: SPAL, ORSD and AG; writing-review and editing: SPAL, ORSD, RJS, BM, AJM and AG. Supervision: BM, AJM and AG; funding acquisition: BM, AJM, RJS and AG.

## DECLARATION OF INTEREST

The authors declare that they have no conflicts of interest related to the contents of this article.

## SUPPLEMENTAL ITEMS

**Fig S1.**
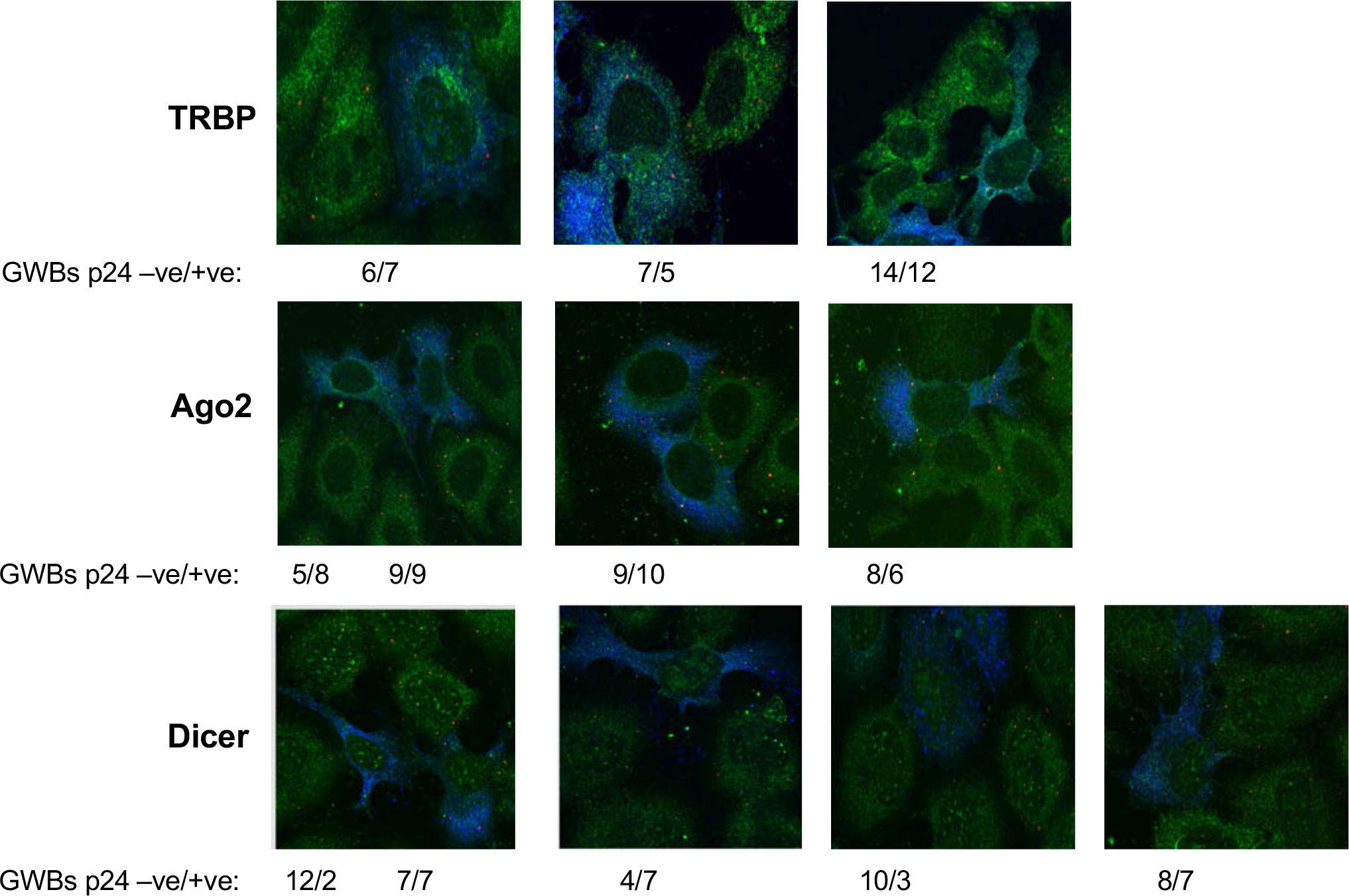
GWB counts in side-by-side images of p24 +ve and –ve cells. HeLa cells were transfected with 0.2 μg of HIV-1 pNL4-3. 48 h later cells were fixed and stained with the human GWB marker serum 18033 (red), an antibody targeting HIV-1 p24 (blue) and an antibody targeting TRBP, Ago2 or Dicer (green). The level of red was adjusted with Adobe Photoshop so that only large GWBs (2-14 per cell) were visible. GWBs were then counted in side-by-side images of p24 –ve/+ve cells and recorded below the images.

**Fig S2.**
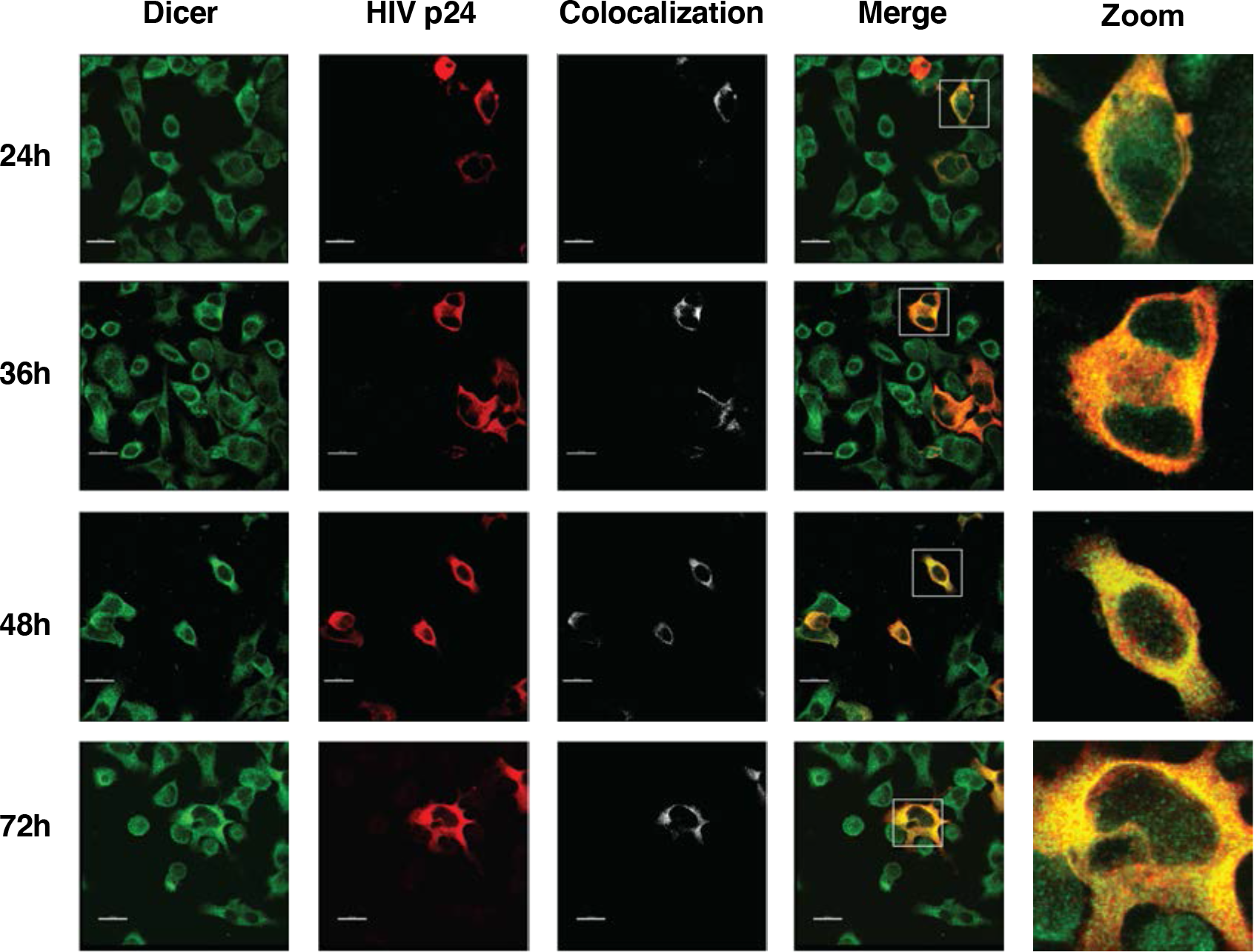
Viral Gag protein colocalizes with Dicer in HIV-1 producing cells. HeLa cells were transfected with 0.2 μg of HIV-1 pNL4-3. Cells were subjected to transfection time points of 24 h, 36 h, 48 h and 72 h. After each time point, cells were fixed and stained with mouse anti-Dicer 13D6 (green) and rabbit anti-p24 (red). The size scale is 20 μm and is shown in each picture on the bottom left. Digitally zoomed images of the merged channels are shown on the far right. A colocalization channel was built using Imaris software and is displayed in the third lane. The calculated Pearson’s correlation coefficient for each time point is the average from 5 cells ±SEM and is: 0.5420 ± 0.0532 at 24 h; 0.5320 ± 0.0258 at 36 h; 0.6380 ± 0.0290 at 48 h; 0.5560 ± 0.0660 at 72 h.

**Fig S3.**
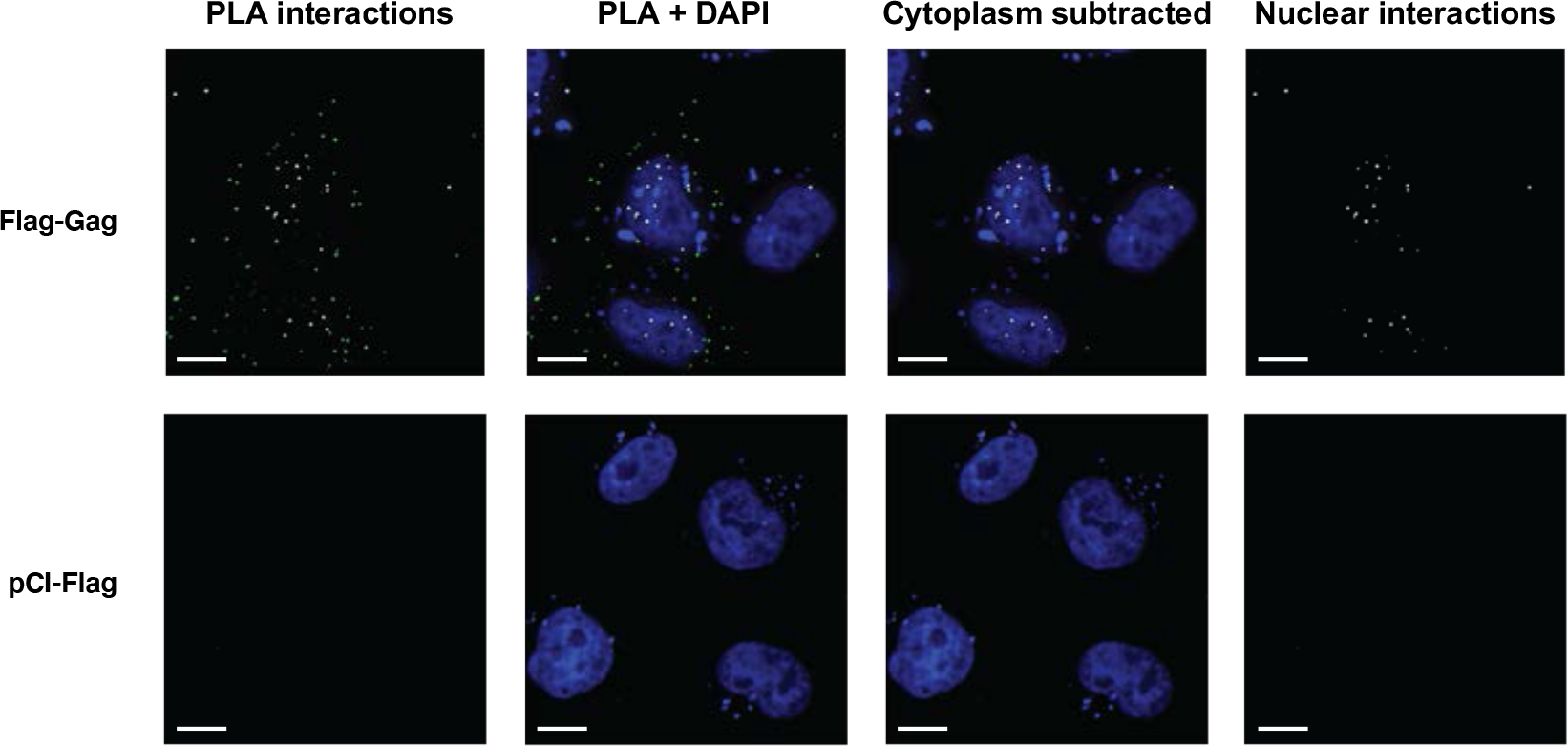
Dicer and Gag PLA. Images of transfected HeLa cells with pFlag (bottom) or Flag-Gag (top) are representative of 100 counted cells. The size scale is 10 μm and is shown in each picture on the bottom left. From left to right: the first column shows a representative field in the PLA channel. The second column shows the merged channel of PLA and DAPI. The third column shows the merged channel, only including PLA signals within the nucleus. The fourth column shows the colocalization signal between PLA and DAPI (interactions within the nucleus).

**Fig S4.**
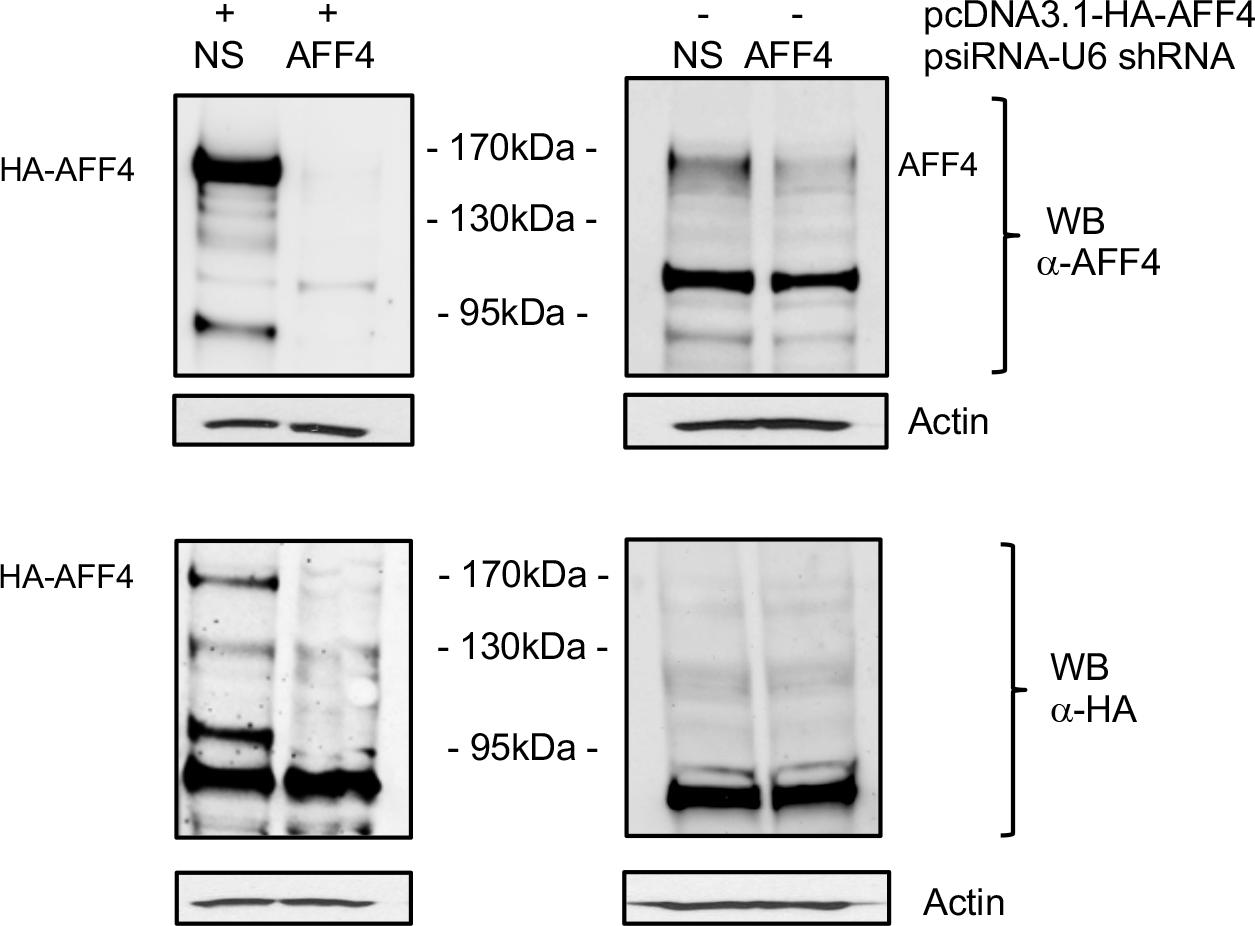
Validation of AFF4 antibody. HEK 293T cells were co-transfected with pcDNA3h/TO-HA-AFF4 or no plasmid (endogenous AFF4) in addition to either psiRNA-U6GFP::Zeo-shNS or psiRNA-U6GFP::Zeo-shAFF4. 30 μg (left part with transfected HA-AFF4) or 100μg (right part showing endogenous AFF4) of protein extracts were separated on two identical 10% SDS-PAGE gels and analyzed by Western blot using anti-AFF4 (7D) or anti-HA and anti-actin as indicated. The upper right part of the blot is identical to the left part of Figure 6D because the pictures originate from the same gel and the same blot. The blots shown are representative of three independent experiments.

**Supplementary table 1.** Specific RNAs in Dicer during the occupancy analysis.

**Supplementary table 2.** Specific RNAs in Dicer-Gag during the occupancy analysis.

**Supplementary table 3.** RNAs found in Dicer and Dicer-Gag during the occupancy analysis.

**Supplementary table 4.** Normalized coverage of consensus and confirmed miRNA sequences.

**Supplementary table 5** TargetScan predicted human miRNA regulators of AFF4 mRNA (ENST00000265343.5) 3’UTRs, sorted by cumulative weighted context score.

